# Removal of nascent transcripts by TTF2 is required for sister chromatid resolution in human cells

**DOI:** 10.64898/2026.01.26.701825

**Authors:** Laura Tovini, Inês Milagre, Catarina Peneda, Raquel A. Oliveira

## Abstract

During mitosis, chromosome assembly is accompanied by a global shutdown of transcription. However, how this transcriptional silencing contributes to mitotic fidelity and genome stability remains poorly understood. Here, we used depletion of Transcription Termination Factor 2 (TTF2) – a key factor in mitotic transcriptional inactivation – to investigate the impact of pervasive transcription on mitotic fidelity. TTF2 depletion causes accumulation of elongating transcripts on mitotic chromatin and multiple mitotic defects, including abnormal chromosome alignment, delayed progression, and impaired chromosome compaction. Notably, defects in sister chromatid resolution are particularly prominent, with DNA bridges as the major segregation error, increasing micronuclei formation. These defects are linked to altered chromatin organisation, including R-loops accumulation at mis-segregating DNA. Most anaphase defects are suppressed when transcription is chemically inhibited, establishing a causal link between transcription and the observed mitotic defects. Our findings reveal how abnormal retention of transcriptional activity on mitotic chromatin disrupts mitosis, with impaired sister chromatid resolution linking transcriptional dysregulation to genome instability.

## Introduction

Transcriptional silencing at mitotic entry is a conserved feature of metazoan cells. While this process has long been believed to be a secondary effect of chromosome condensation, an increasing number of studies suggest that active mechanisms are in place to ensure timely transcriptional shut-off at mitotic entry (Contreras and Perea-Resa, 2024). Possible functions of mitotic transcriptional silencing include limiting the transcription retention to the centromeric regions to allocate key centromere/kinetochore factors (Perea-Resa et al., 2020; Blower, 2016; McNulty et al., 2017), ensuring proper chromosome condensation (Ramos-Alonso et al., 2023; Lebreton et al., 2024), or favouring differential gene expression reprogramming (Wojenski et al., 2021; Pelham-Webb et al., 2021; Palozola et al., 2017). Despite these ideas, analysing the consequences of faulty transcriptional silencing on mitotic fidelity remains challenging due to the limited knowledge of the factors involved.

One player that has emerged in the termination of RNA polymerase II transcription is Lodestar (Lds)/Transcription Termination Factor 2 (TTF2), whose depletion leads to the retention of nascent transcripts in both mammalian cells and *Drosophila* embryos (Jiang et al., 2004; Carmo et al., 2023). Lds/TTF2 is an ATPase that lies in the cytoplasm during interphase and translocates on chromatin only upon nuclear envelope breakdown (Carmo et al., 2023; Fujisawa & Labib, 2024; Jiang et al., 2004). *In vitro* experiments showed that Lds/TTF2 is responsible for the displacement of RNA polymerase II and its transcripts from a DNA template, independently of RNA length and regardless of the modification state of the polymerase (Liu et al., 1998; Jiang et al., 2004). Although Lds/TTF2 depletion has been associated with abnormal mitotic divisions (Jiang et al., 2004; Carmo et al., 2023), the mechanistic basis of the resulting chromosome segregation defects—particularly their causal link to transcriptional persistence on mitotic chromosomes—has remained unresolved. Depletion of Lds in Drosophila embryos led to severe mitotic failures that could not be reverted by transcription inhibition (Carmo et al., 2023). It was therefore suggested that Lds additionally participates in the resolution of sister chromatid entanglements, possibly by promoting topoisomerase II activity (Carmo et al., 2023). Additionally, recent studies in mammalian cells revealed that TTF2 is involved in the displacement of DNA polymerases from the mitotic chromatin of cells under mild replication stress (Fujisawa and Labib, 2024; Can et al., 2024). Hence, the exact contribution of nascent transcript retention, caused by Lds/TTF2 depletion, on mitotic fidelity remains unclear.

Here, we use TTF2 knockdown to induce the retention of transcripts on mitotic chromatin in HeLa cells and probed for the consequences on mitotic fidelity. Our results indicate that mitotic transcriptional silencing is required for the efficiency of several mitotic pathways and the preservation of genome integrity, which, if perturbed, may predispose cells to cancer and/or other human diseases linked to genomic instability.

## Results

### TTF2 KD induces transcript retention on mitotic chromosomes

To determine the importance of transcripts eviction in mitosis for faithful cell division, we used TTF2 knockdown as a tool to impair mitotic transcriptional silencing in HeLa cells. TTF2 has been previously reported to be involved in the removal of engaged RNA polymerase II from metaphase chromosomes (Carmo et al., 2023; Jiang et al., 2004) To confirm these observations, we treated HeLa cells with a control siRNA or TTF2-specific siRNA for 48 hours, which allows for efficient depletion of TTF2 (**Supplementary Fig. 1a**). To assess the degree of transcript retention on mitotic chromatin, we incubated cells with 5-Ethynyl Uridine (EU) for 30 min to be able to detect nascent mitotic transcripts. Upon fixation and ClickIT chemistry we analysed the retention of nascent transcripts by measuring the enrichment of EU (mean value on the DNA area relative to the mean value of the whole cell area), in interphase cells and in each mitotic phase (**Fig. 1a-d**).

**Figure 1.**
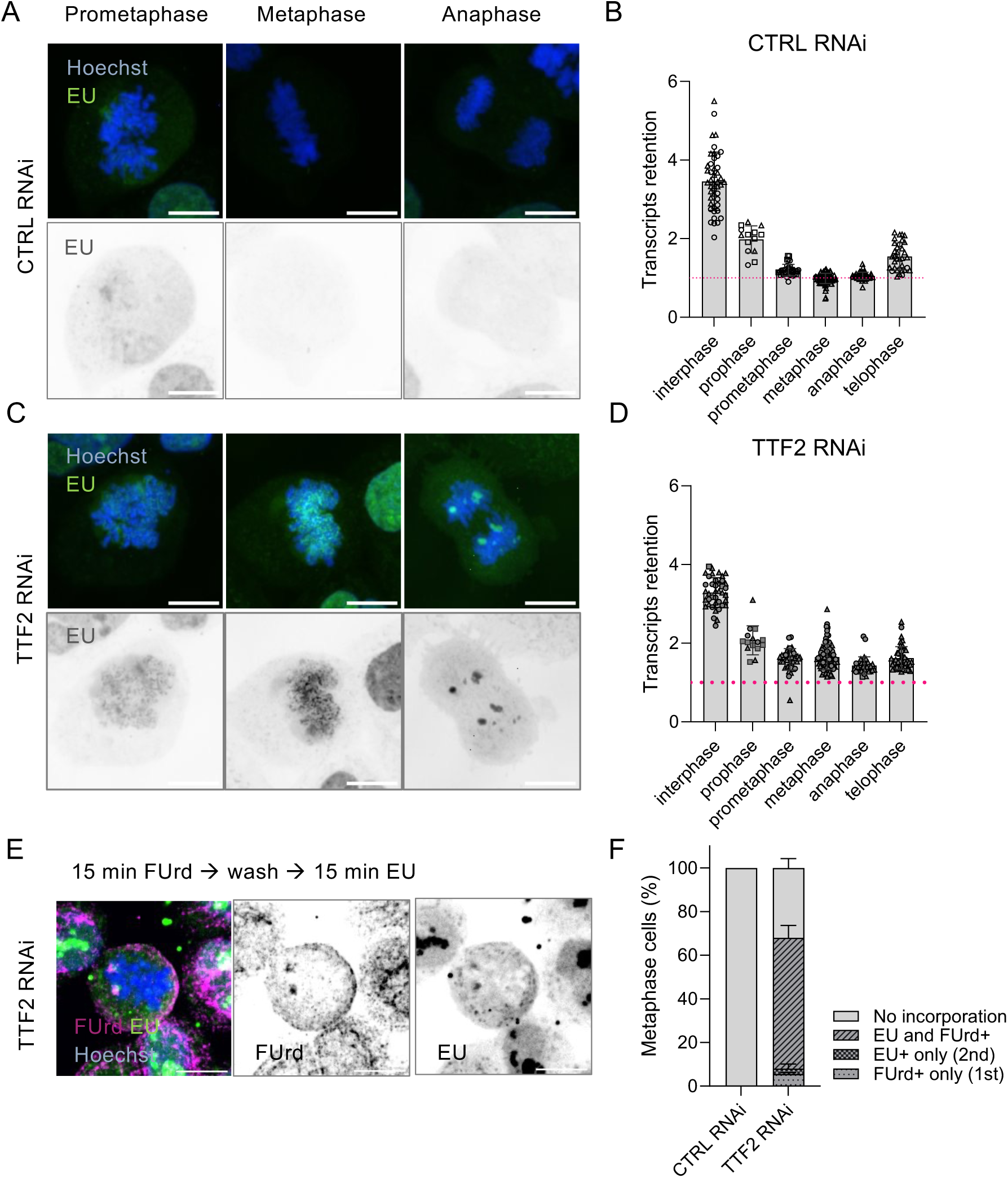
TTF2 KD induces transcript retention and continuous RNA pol II elongation in mitosis. **A,C**) Representative images of HeLa mitotic stages upon CTRL/TTF2 KD and EU incorporation. DNA is in blue (Hoechst) and nascent transcripts are in green (EU). **B,D)** EU signal retention on DNA area relative to whole cell area in interphase and in each mitotic phase indicated. Each dot represents one cell, with bars indicating the mean value of 3 independent experiments. Pink dotted line indicate the baseline EU retention in control mitotic cells (=1). **E**) Representative images of HeLa metaphases upon TTF2 KD and Uridine-analogues sequential incorporation (EU in green, FUrd in magenta, DNA in blue.) **G**) Quantification of metaphase cells (%) that incorporated the first analogue only (FUrd+), the second analogue only (EU+), or both analogues (EU and FUrd +). Data are from 2 independent experiments, bars indicate the mean and SD. Scale bars are all 10 μm.

Control siRNA-treated cells exhibit detectable chromatin enrichment of EU-signals early in prometaphase, but as cells progress through mitosis, the amount of EU that gets retained on mitotic chromatin decreases and reaches minimal values in metaphase and anaphase (retention index = 1, i.e., no enrichment on chromatin, pink dotted line in **Fig. 1b**), stages at which the transcription appears fully silenced. Later, as cells reach telophase and chromosomes start to decondense, the transcriptional process starts again, as visible by an increased amount of EU retention on chromatin (**Fig. 1a**,**b**).

In contrast, cells treated with TTF2 siRNA showed a similar enrichment of interphase transcripts after mitotic entry (prometaphase), but a higher retention in the subsequent phases of mitosis (60% enrichment in metaphase and 40% enrichment in anaphase compared to the control). Thus, TTF2 KD mitotic cells showed a consistent increased accumulation of EU-labelled transcripts across all stages of mitosis compared to control cells. The most striking phenotype was observed in metaphase, where EU-labelled transcripts colocalise all over mitotic chromatin (**Fig. 1c**, central panel).

To rule out the presence of off-target effects, a TTF2 siRNA-resistant cell line was generated by introducing silent nucleotide substitutions (Ong, 2021) within the siRNA target site(s) without altering the encoded amino acid sequence (**Supplementary Fig. 1)**. We observed that, in contrast to the siRNA-sensitive and the parental cell lines, the siRNA-resistant line showed no detectable decrease of TTF2 (**Supplementary Fig. 1a**) and only 20% of cells retained EU on chromatin in metaphase, compared to nearly 100% in the sensitive and parental lines (**Supplementary Fig 1b**). To confirm that the retained transcripts originate from RNA polymerase II (i.e., mRNAs), we tested whether transcript retention observed in TTF2 KD mitotic cells could be reverted by RNA polymerase II inhibition. Cells were treated with Triptolide (TRP) for 4 hrs (Titov et al., 2011), and retention of nascent transcripts was evaluated as above. Quantitative analysis of nascent RNAs on mitotic chromatin revealed that EU levels on chromatin were reduced to CTRL values (**Supplementary Fig. 2**), confirming previous observations that TTF2 depletion results in mRNAs retention in mitosis.

Taken together, these data reveal that TTF2 is involved in silencing transcription upon mitotic entry, driving the removal of RNA polymerase II-dependent transcripts that otherwise get retained during the entire length of mitosis.

### Transcripts retained by TTF2 KD continue elongation on mitotic chromatin

The experiments described above use a 30 min EU-incorporation time, which is only slightly lower than the mitotic duration in these cells (NEBD to anaphase onset 43.3 min +/- 7.5 min, median values). Moreover, even in control cells, a significant amount of nascent transcripts is observed on chromatin of prometaphase cells (**Fig. 1a,b**). Therefore, it is unclear if the transcripts detected on mitotic chromatin in TTF2-depleted cells correspond to RNAs transcribed in late G2/early mitotic stages that we have not been efficiently evicted and/or these transcripts are the result of continued transcription elongation after NEBD. Previous studies suggested the latter possibility, as cells with reduced TTF2 levels exhibited a dramatic retention of Ser2 phosphorylated Pol II (“active RNA PolII”) on mitotic chromosomes (Jiang et al., 2004; Fujisawa and Labib, 2024). Nevertheless, it is still possible that RNA PolII remains in an active biochemical state, yet efficient elongation is prevented by the changes that occur in mitotic chromatin. To test whether retained transcripts can continue elongation during mitosis, we used a shorter and sequential dual labelling protocol (15 min of 5-FluoroUridine, FUrd, in the first step of incorporation, followed by 15 min of EU) so that we can differentiate transcriptionally stalled mitotic cells, which will retain only the first analogue (FUrd), from transcriptionally active mitotic cells, which will efficiently incorporate both analogues (FUrd and EU) (**Fig. 1e**). This analysis revealed that in the absence of TTF2, the transcripts retained on most metaphase cells (∼60%) showed incorporation of both analogues (**Fig. 1f**). Control experiments demonstrate that neither staining detects the other analogue, confirming specificity and absence of cross-reactivity (**Supplementary Fig. 3**). Considering the short period of the second analogue incorporation (15 min), way below the whole mitotic duration, we infer that the transcripts labelled with this analogue result from active elongation during mitosis.

These experiments reveal that in the absence of TTF2, at least some transcripts continue to be actively elongated on mitotic chromosomes.

### TTF2 depletion disrupts metaphase organisation

Our results thus far indicate that in the absence of TTF2, actively elongating transcripts accumulate on mitotic chromatin. We next sought to investigate whether a transcriptionally permissive chromatin environment, together with the retained transcripts, could potentially impact mitosis. To monitor the mitotic progression over time, we recorded HeLa cells in control or TTF2-depleted conditions by time-lapse imaging. We observed that in metaphase cells—defined here as the frame acquired 3 minutes (one time-point) before anaphase onset (AO)—chromosomes were spread over a wider area (**Fig. 2a,b**). These findings suggest that the presence of nascent transcripts on mitotic chromatin disrupts metaphase organisation.

**Figure 2.**
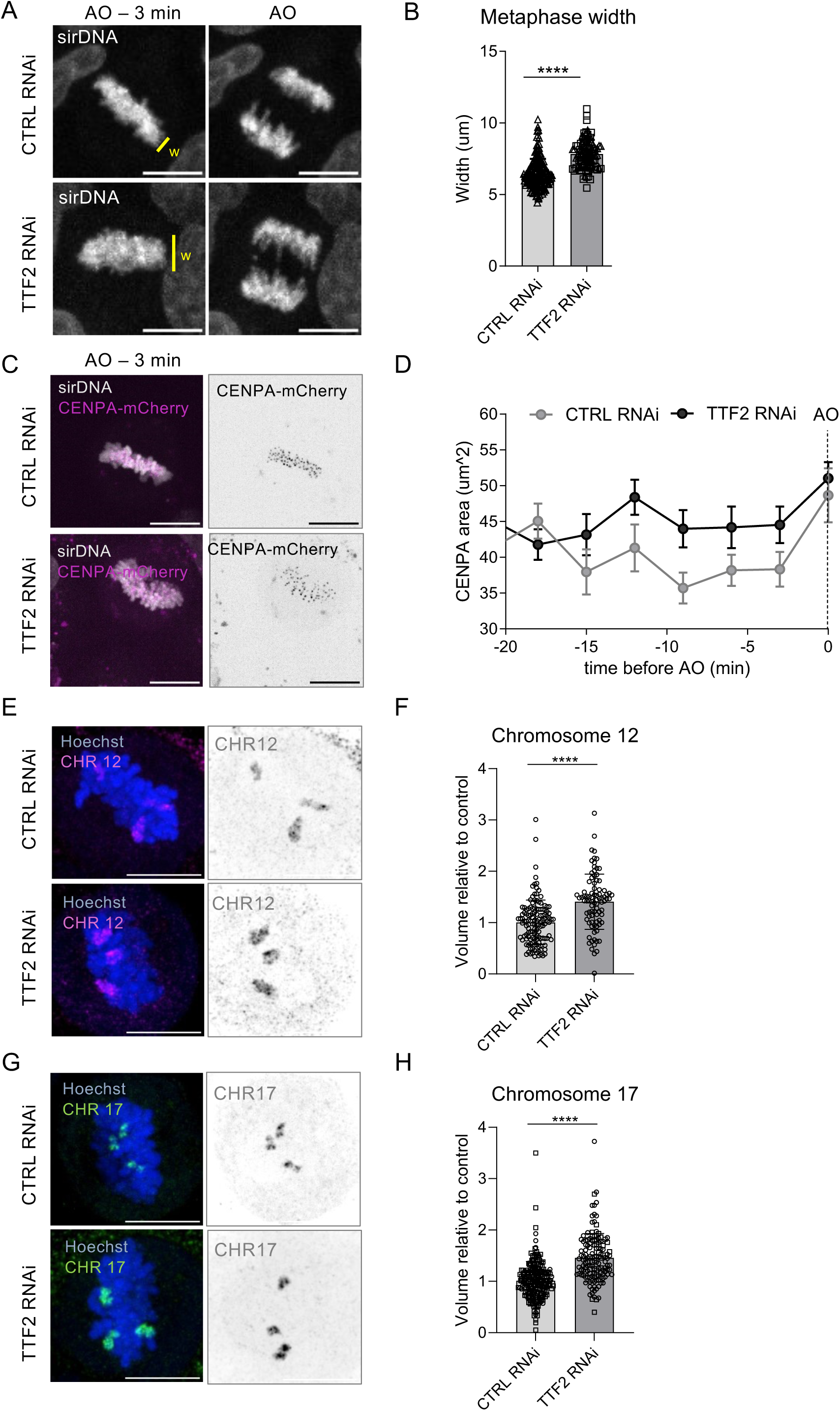
TTF2 depletion disrupts metaphase organisation. **A)** Snapshot of CTRL/TTF2 KD cells from live-cell imaging, one frame before AO (-3 min) and at anaphase. DNA is in grey and the yellow line indicates the measurements performed in B. **B**) Measurements of one frame before AO. Each dot represents a cell, bars indicate mean and SD. Data are from 3 independent experiments. **C**) Representative images of metaphase cells in CTRL/TTF2 KD HeLa-mCherry one frame before AO. DNA is in grey, CENPA is in magenta. **D**) Measurements of the whole area occupied by CENPA dots in live-cell imaging during mitosis up to AO. Each dot represents the mean value of measurements collected with 3 independent experiments. **E,G**) Representative images of whole chromosome FISH (chromosome 12 and 17) on CTRL (top) and TTF2 (bottom) KD HeLa. **F,H**). Quantification of single chromosomes volume normalised to control condition. Each dots represents one chromosome, bars indicate the mean and SD. Scale bars are all 10 μm. ****p<0.0001. *p*-values were calculated using Student’s t-test.

Further analysis on chromosome alignment, using a CENPA-mCherry–tagged cell line (a gift from the Kops lab), revealed that upon TTF2 depletion, centromeres occupy a larger area compared to controls. This increase is observed throughout prometaphase and metaphase, including at the final metaphase time point immediately preceding anaphase onset (**Fig. 2c**, -3 min relative to AO, and **Fig. 2d**). This analysis indicates that TTF2 depletion causes mild alignment problems, which remain uncorrected prior to anaphase onset.

Accordingly, by plotting the time that each cell spends from NEBD to AO, we found a modest but significant Spindle Assembly Checkpoint (SAC)-dependent delay in TTF2 KD condition, with an increased average time that goes from 36.2 +/- 7.4 min in control to 43.3 +/- 7.5 min in TTF2 KD (**Supplementary Fig. 4a,b**). Despite this delay, TTF2-depleted cells do not show evidence of the canonical triggers for SAC activation: 1) the percentage of fully uncongressed chromosomes is not significantly higher than in control conditions (**Supplementary Fig. 4d,e**); 2) the inter-centromere distance before AO (-3 min), used as a proxy for kinetochore–microtubule (KT-MT) attachment stability (Jennifer C Waters et al., 1996), showed no difference between conditions (**Supplementary Fig. 4f,g**) 3) the number of Mad2-positive kinetochores in prometaphase/metaphase cells also showed similar distributions across both conditions (**Supplementary Fig. 4h,i**).

Together, these observations suggest that chromosomes in TTF2-depleted cells successfully establish bipolar attachments, yet their alignment at the metaphase plate is slightly impaired by mechanisms independent of KT–MT attachment. Because accurate metaphase organisation reflects both kinetochore function and chromosome-intrinsic properties, we next examined chromosome assembly and chromatin organisation. To evaluate the level of chromosome compaction, we performed chromosome-specific fluorescence *in-situ* hybridization (FISH) and quantified the volume of individual chromosomes in control and TTF2-depleted metaphase cells. For both chromosomes examined (chromosome 12 and chromosome 17), TTF2 depletion resulted in an approximately 1.5-fold increase in chromosome volume relative to controls (**Fig. 2e-h**). These results indicate that retention of nascent transcripts on mitotic chromosomes impairs chromosome assembly.

Collectively, these findings demonstrate that TTF2 depletion leads to a spectrum of metaphase defects, including altered chromosome organization, reduced alignment precision, and delayed anaphase onset, while preserving bipolar attachment and tension.

### Transcript retention on mitotic chromatin leads to DNA bridges in anaphase

Given the broad spectrum of metaphase defects observed upon TTF2 depletion, we next examined the fidelity of anaphase chromosome segregation to determine whether mitotic accuracy is affected. Based on the live-cell imaging data, we quantified chromosome segregation errors and classified them into two categories (**Fig. 3a** top row and **b**): *laggings* – when a single chromatid or chromatin fragment fails to move efficiently towards either anaphase side, or *bridges* – when a continuous thread of DNA is connecting the two sides of anaphase. This analysis revealed that anaphase cells depleted of TTF2 displayed significantly higher chromosome segregation rates compared to the control (approximately 30% more), with anaphase bridges representing the most predominant defect (**Fig. 3a,b**). The frequency of chromosome laggings was modest (9%), not statistically different from controls (∼3%). Consistently, analysis of chromosome mis-segregation in fixed cells revealed a similar phenotype, with TTF2-depleted cells displaying on average 18.6% increase in anaphase bridges relative to control cells, with few laggings observed (1.3%) (**Fig. 3b** bottom row and **c**). Importantly, when cells were treated for a short period of time with triptolide (TRP; 2h, minimal time sufficient to rescue most of transcript retention in interphase, **Supplementary Fig. 5**), the amount of anaphase bridges was reverted to near control levels (7.6% in average, **Fig. 3c)**. We therefore conclude that chromosome segregation upon TTF2 KD is compromised by transcription-related events, with DNA bridges emerging as the main observed defects. Live-cell imaging allowed us to follow the fate of chromosome mis-segregation events in G1. A small but significant fraction of dividing TTF2-depleted cells resulted in micronuclei at mitotic exit (∼12% compared to 1.3% in the control). These formed either around lagging chromosomes/chromatin fragments (5.6%) or from DNA bridges (6.3%) (**Fig. 3d,e**).

**Figure 3.**
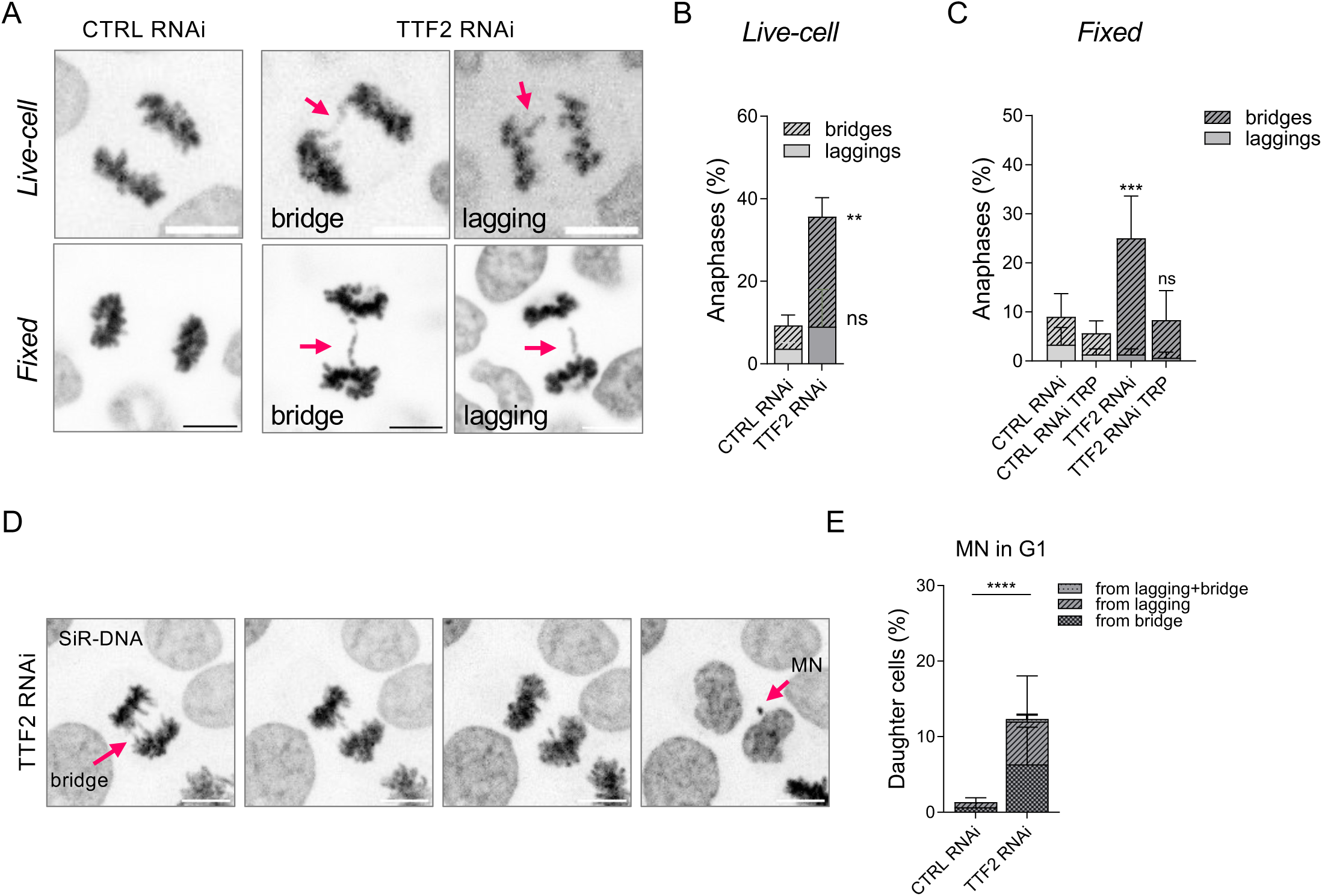
Transcripts retention on mitotic chromatin leads to DNA bridges in anaphase and MN. **A**) Representative images of control anaphases and defects commonly seen in TTF2 KD HeLa cells from fixed and live-cell imaging. **B**) Frequency of anaphase cells with the chromosome mis-segregation defects indicated in CTRL/TTF2 KD HeLa cells from live-cell imaging. Bars represent mean and SD from 3 independent experiments. *p*-value was calculated using a two-way ANOVA with multiple comparisons, **p=0.0016. **C**) Frequency of anaphase cells with the chromosome mis-segregation defects indicated in CTRL/TTF2 KD, DMSO or Triptolide (TRP) treated HeLa cells from fixed imaging. Data represent mean and SEM from 3 independent experiments. *p*-value was calculated using a two-way ANOVA with multiple comparisons, ***p=0.0007. **D**) Example images of HeLa cells exiting mitosis upon TTF2 RNAi. Arrows indicate a lagging chromosome that is going to form a micronucleus (MN) in the daughter cell. **E**) Frequency of cells (%) with MN and their origin as indicated in the legend. Data comes from 3 independent experiments. Bars represent mean and SD. *p*-value was calculated using an Ordinary one-way ANOVA (p<0.0001) for comparison of the whole amount of MN. Scale bars are all 10 μm.

Considering that anaphase bridges are usually associated with unresolved DNA threads (Finardi et al., 2020), we tested whether additional fibres that cannot be detected with common dyes (ultrafine bridges, UBFs) might also be contributing to the observed phenotype. To do so, we stained UBFs by immunofluorescence with RPA70 and PICH antibodies, which detect ssDNA and dsDNA, respectively (Sarlós et al., 2018). We first scored the percentage of cells carrying any type of UFBs (PICH or RPA70), with or without accompanying DNA bridges (here referred to as DAPI bridges) (**Fig. 4a,b**). While a fraction of UFBs is prevalent also in unperturbed mitotic cells, representing the last attempt of sister chromatids resolution at anaphase onset (Biebricher et al., 2013; Chanboonyasitt and Chan, 2021), we observed an increase of nearly 20% in TTF2 KD cells compared to the control, where the most predominant category was UFBs+ DAPI+ bridges (**Fig. 4b**). To assess the severity of the phenotype, we next quantified the number and type of UFBs per anaphase cell (**Fig. 4c-e**). In control cells, the majority of PICH-positive UFB-containing anaphases displayed either zero or a single fibre. In contrast, TTF2-depleted cells showed a marked increase, with over 40% of anaphases exhibiting one fibre and 36% displaying two or more fibres per cell. Notably, these values returned to near-control levels following a 2-hour triptolide treatment (**Fig. 4d)**. In contrast, we did not detect a major increase in RPA70-coated UFBs (commonly originating from other anomalies, such as DNA replication stress (Chan et al., 2018) upon TTF2 depletion. In ∼20 % of TTF2-depleted anaphases, we observed a single RPA70+ fibre, a frequency that was reduced to nearly zero upon triptolide treatment (**Fig. 4e**). Collectively, these results indicate that TTF2 depletion leads to an increase in transcription-dependent UFB, particularly PICH-positive, suggesting the presence of unresolved sister chromatid intertwines.

**Figure 4.**
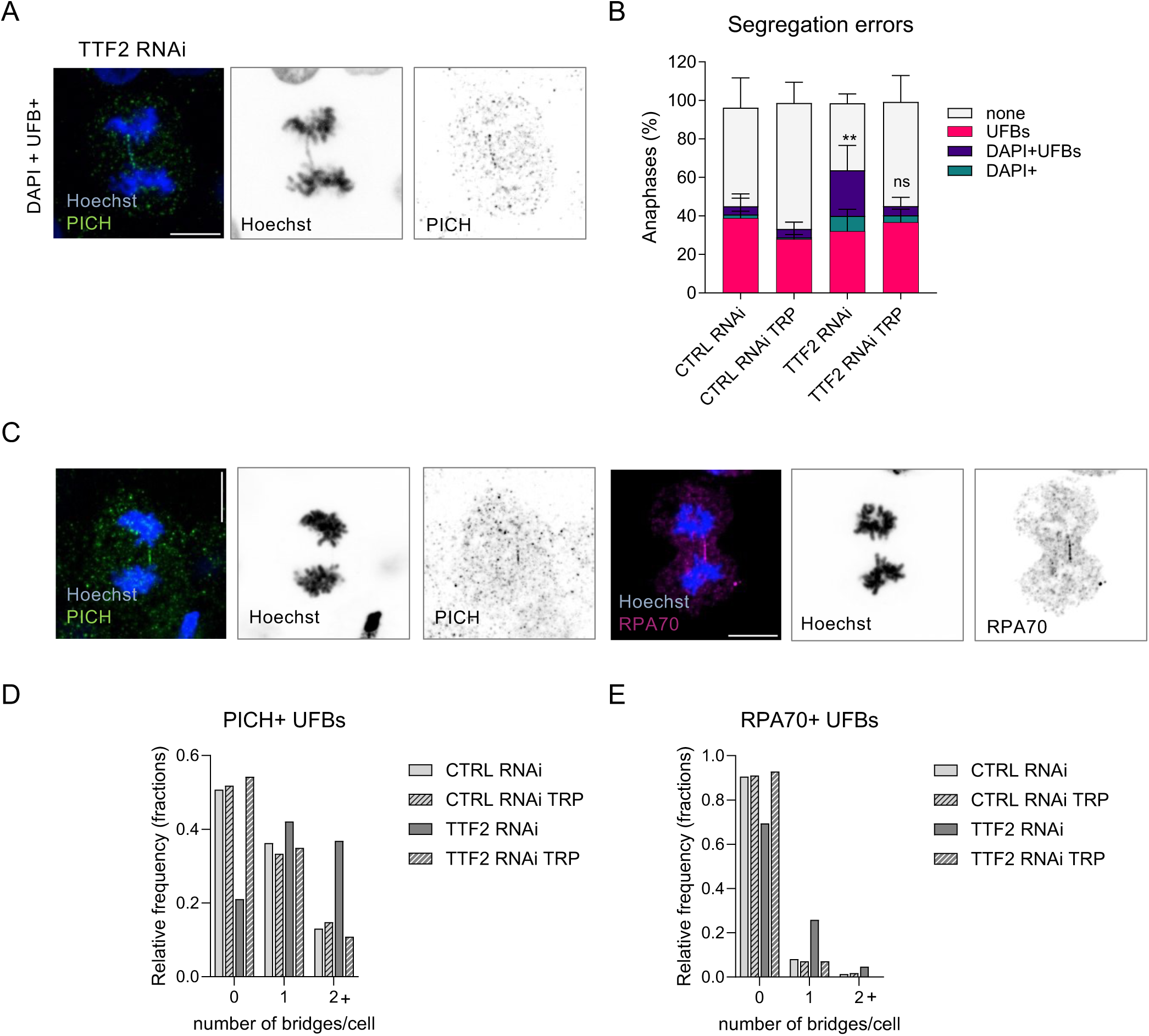
Transcripts retention on mitotic chromatin leads to DNA bridges and UFBs in anaphase and can be reverted upon short RNA pol II inhibition. **A**) Representative images of a TTF2 KD anaphase cell with the most common defect: DAPI+ PICH+ ultrafine anaphase bridges (PICH in green, DNA in blue). **B**) Frequency of anaphase cells with UFBs of each category indicated, where DMSO/TRP treatment was 2h. Data represents mean and SEM from 3 independent experiments. Statistical analysis was performed using two-way ANOVA across four conditions for each variable, followed by Tukey’s multiple-comparison test. Significant differences between conditions are indicated, where **p=0.0015. **C**) Representative examples of the two classes of UFBs analysed in TTF2 KD anaphase cells: PICH (on the left, in green) and RPA70 (on the right, in magenta). **D**) Number of PICH+ UFBs (frequency per cell) in CTRL/TTF2 RNAi and with 2h DMSO/TRP treatment. **E)** Number of RPA70+ UFBs (frequency per cell) in CTRL/TTF2 RNAi and with 2h DMSO/TRP treatment. Bars indicate the mean of 3 independent experiments. Scale bars are all 10 μm.

UFBs are also known to delay sister chromatid separation during anaphase, as entangled chromatin can counteract spindle pulling forces. To test the efficiency of chromatid separation at anaphase onset, we performed live-cell imaging at higher temporal resolution (1 min timeframe) and quantified the distance travelled by segregating chromatids, up to 5 min post-AO (**Supplementary Fig. 6**). As a positive control, we treated cells with mild doses of ICRF-193 (a Topoisomerase IIα inhibitor (Huang et al., 2001) that prevents decatenation and therefore is known to delay sister chromatid separation at anaphase onset (Oliveira et al., 2010). While in DMSO-treated control cells, sister chromatids move at 2.14 μm/min, ICRF-treated cells displayed a less efficient sister chromatid separation (1.54 μm/min). Strikingly, TTF2-depleted cells exhibited a similar defect (1.44 μm/min), supporting the notion that impaired sister chromatid resolution underlies the segregation defects observed upon TTF2 loss (**Supplementary Fig. 6c**). We therefore concluded that anaphase velocities upon TTF2 depletion are consistent with the presence of unresolved intertwines.

Together, these findings indicate that TTF2 depletion sensitises cells to chromosome segregation defects, particularly the formation of anaphase (ultrafine)bridges and micronuclei.

### TTF2 depletion does not impair global topoisomerase 2 localisation or enzymatic activity

Given the high frequency of DNA bridges observed upon TTF2 depletion, we next evaluated the levels and activity of the major enzyme that resolves sister chromatid intertwines during mitosis: topoisomerase IIα (Top2A).

We first asked whether the defects seen upon TTF2 KD might originate from a defective loading of Top2A on mitotic chromosomes, thereby limiting its ability to resolve entanglements. To investigate this, we performed pre-extraction-based immunofluorescence using Top2A antibodies, enabling selective visualisation of chromatin-bound Top2A. Quantification of mean chromatin-associated Top2A intensity revealed comparable levels in both interphase and mitotic cells across control and TTF2-depleted conditions, suggesting no major defect in Top2A chromatin-association (**Fig. 5a,b**). However, steady-state binding measurements may not capture differences in enzymatic activity or residence time. To overcome this limitation, we implemented a Differential Retention of Topoisomerase (DRT) assay (Agostinho et al., 2004) in HeLa cells to evaluate the level of Top2A strand passage activity.

**Figure 5.**
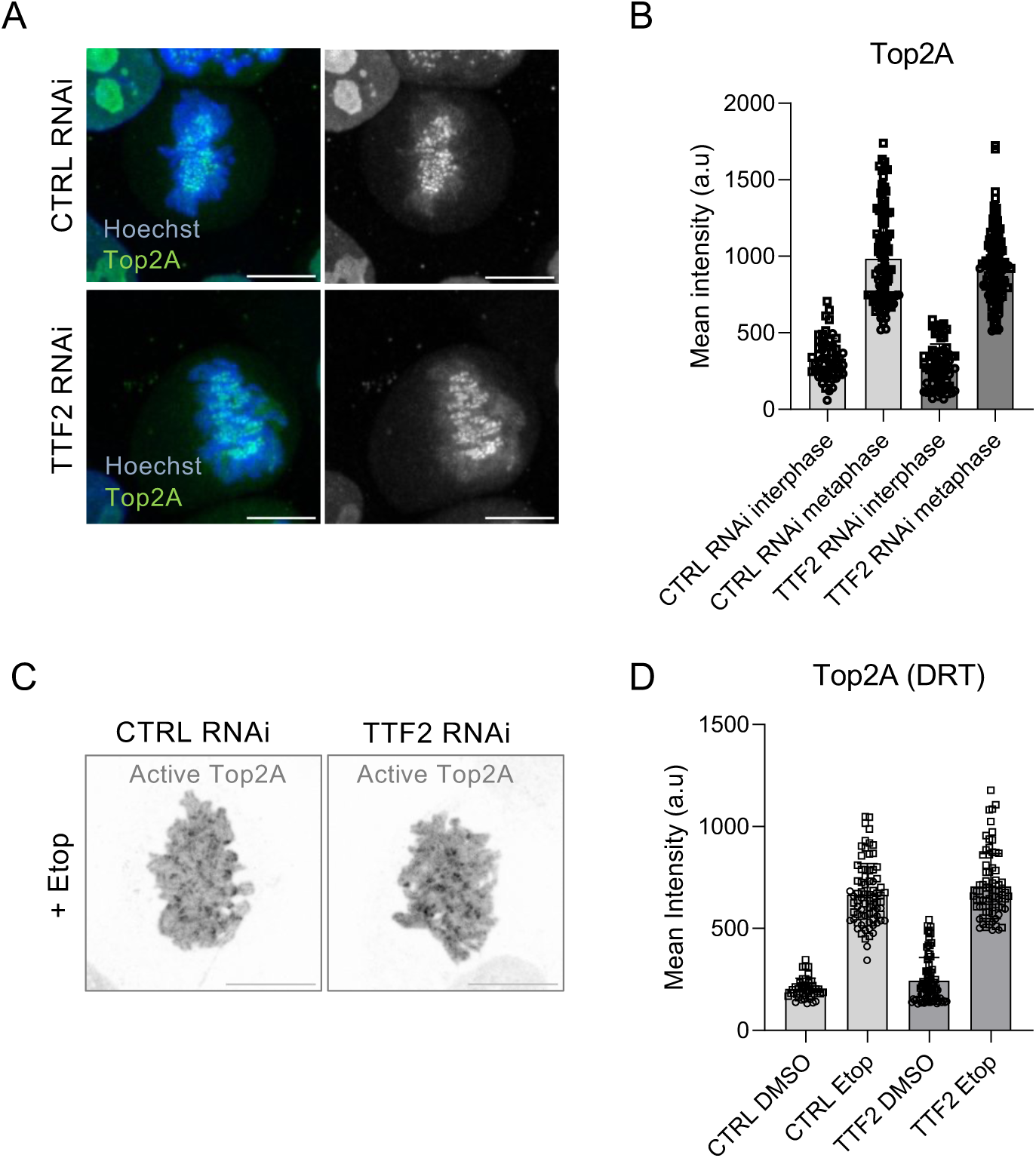
TTF2 does not impair global localisation or enzymatic activity of Top2A. **A**) Representative images of Top2A staining on CTRL/TTF2 KD HeLa cells. Top2A is in green, DNA in blue. **B**) Quantification of Top2A mean intensity levels in metaphase. Each dot represents one cell, bars indicate the mean and SD of 2 independent experiments. **C**) Representative images of Active Top2A staining upon DRT assay (Etop=etoposide) in metaphase cells in CTRL/TTF2 KD HeLa. **D**) Quantification of Active Top2A mean intensity levels in metaphase cells in CTRL/TTF2 KD HeLa. Each dot represents one cell, bars indicate mean and SD from 2 independent experiments. Scale bars are all 10 μm.

In this assay, cells are treated with etoposide, which selectively traps on DNA Top2A molecules that are catalytically engaged in strand passage reactions (Montecucco et al., 2015). These covalently bound molecules are thus resistant to high-salt extraction, which is used to wash away any enzyme that is not entrapped. Subsequent Top2A staining allows for analysis only of the bound and catalytically active Top2A (Active Top2A). The conditions of the original protocol were modified to decrease etoposide concentration and incubation time, to avoid saturation (**Supplementary Fig. 7**). We reasoned that if TTF2 depletion impaired Top2A activity, we would observe reduced Top2A retention on chromatin during metaphase. However, control and TTF2-depleted cells exhibited comparable retention patterns (**Fig. 5c,d**), indicating no detectable defect in Top2A catalytic engagement. We therefore conclude that TTF2 depletion does not impact the global loading or enzymatic activity of Top2A.

### Mitotic transcription upon TTF2 KD causes excessive accumulation of R-loops on mitotic chromatin

The finding that Top2A is efficiently loaded and active prompted us to explore whether DNA bridges are associated with alterations at the substrate level, rather than defects in the molecular players that promote their resolution. Because increased transcriptional activity is commonly associated with elevated DNA:RNA hybrids, we next examined the presence of those structures. To detect DNA:RNA hybrids, we employed S9.6 antibody (Bou-Nader et al., 2022) and assess its binding on (or attached to) the chromatin of CTRL or TTF2 depleted cells. We found that whereas in control conditions DNA:RNA hybrids are nearly absent, upon TTF2 depletion there is a marked increase in the number of S9.6 positive signals within metaphase chromatin area **(Fig. 6a,b**).

**Figure 6.**
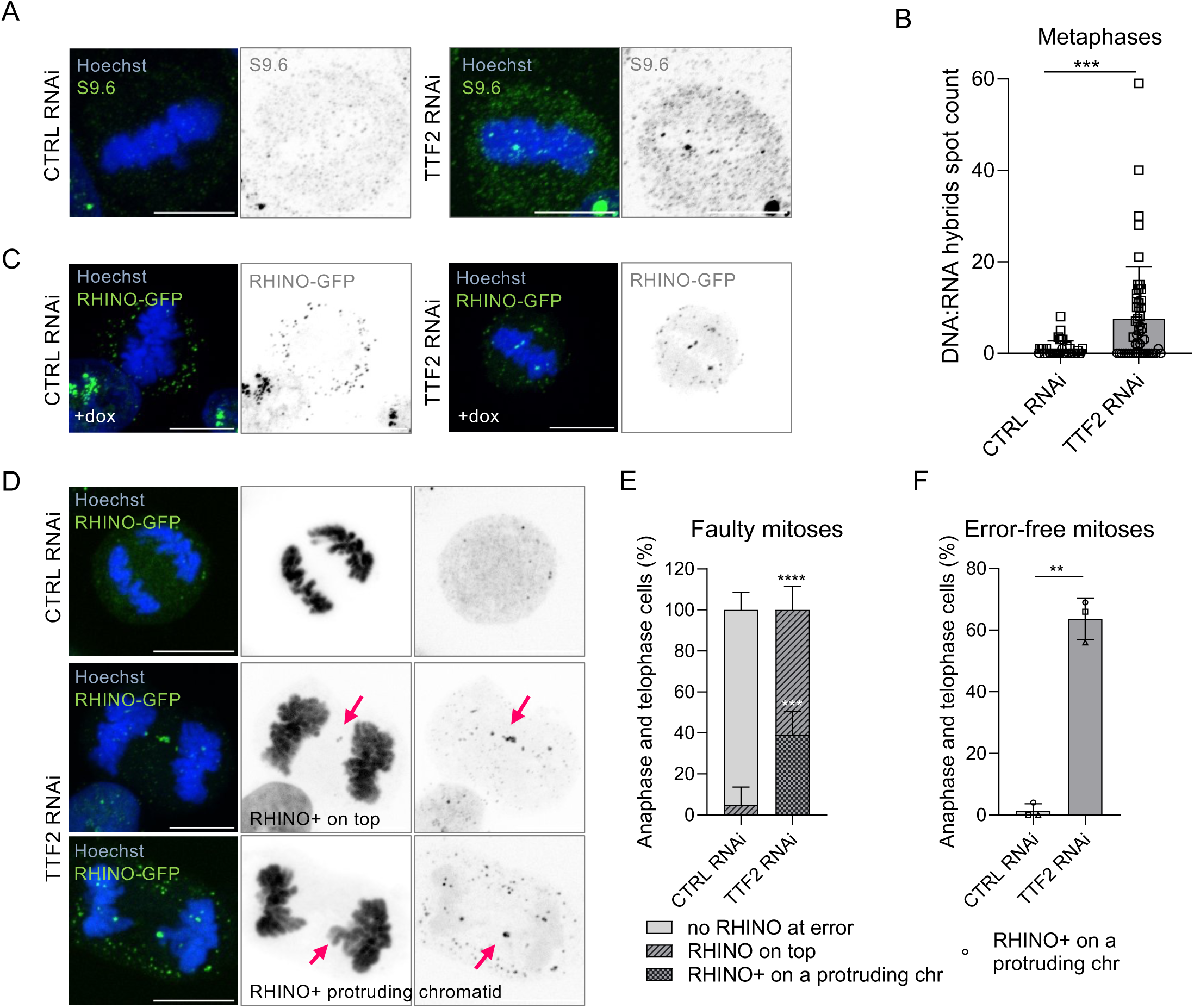
TTF2 depletion causes excessive accumulation of R-loops on mis-segregating chromatids. **A**) Representative images of metaphase cells stained with the S9.6 RNA:DNA hybrid antibody. **B**) Quantification of the number of S9.6 antibody foci detected on DNA in metaphase. Each dot represents one cell, bars indicate the mean and SD of 2 independent experiments. Statistical analysis was performed with a Mann Whitney test. Significant differences between conditions are indicated (*p*<0.05). **C**) Representative images of CTRL/TTF2 KD metaphase HeLa expressing the R-loops sensor RHINO-GFP (in green) after dox induction. **D**) Representative images of CTRL/TTF2 KD HeLa cells with RHINO-GFP in anaphase/telophase. **E**) Frequency of faulty mitoses that do not have a RHINO-GFP signal at the error or show a RHINO-GFP signal either on top of an error or on a protruding chromatid (chr=chromatid). *p*-values were calculated using a one-way ANOVA, ***p=0.0003, ****p<0.0001. **F**) Frequency of error-free mitoses that have a RHINO-GFP+ protruding chromatid. Bars represent mean and SD from 3 independent experiments. *p*-values were calculated using a Student’s *t*-test, **p=0.0026. Scale bars are all 10 μm.

To complement these observations, we also employed RHINO, a genetically encoded RNA:DNA hybrid sensor (Martin et al., 2023), that allows *in situ* visualisation and quantification of R-loops with high specificity, addressing specificity limitations of antibody-based methods. We established a stable HeLa cell line expressing GFP-tagged RHINO upon doxycycline induction. Similar to the previous observation, we found that TTF2 KD metaphases but not the respective control, displayed well defined RHINO foci colocalising with chromatin (**Fig. 6c**), indicating accumulation of bona fide R-loops. To assess the contribution of R-loops to the observed mitotic errors, we evaluated anaphase/telophase cells with detectable mitotic defects for the presence of a RHINO-GFP signal on or near the mis-segregating DNA (**Fig. 5d-f**). In control cells the infrequent mitotic errors (16%) were mostly without any associated RHINO signal. In TTF2 depleted cells, among their faulty anaphase/telophase figures (43%), 61% had RHINO signal on top of the DNA bridge/fragment and the remaining 39% had detectable RHINO foci on chromatids protruding from the segregating mass (**Fig. 6d,e**). These findings indicate that most mis-segregation events are associated with abnormal R-loops retention. Moreover, even in the absence of detectable DNA bridges/fragments, most (63%) of the anaphase/telophase figures in TTF2 depleted cells showed RHINO signal on protruding chromatids (**Fig. 6d,f**).

Together, these results indicate that TTF2-depleted cells display defective chromatin organisation, with abnormal R-loops accumulation that coincide with the sites of impaired segregation.

## Discussion

Here we provide a full characterisation of how abnormal retention of nascent transcripts on mitotic chromatin, upon TTF2 depletion, impacts the fidelity of nuclear division. We further show that retained transcripts are not merely trapped on mitotic chromatin but instead engage in active transcriptional elongation. These findings indicate that failures in transcriptional termination not only may impose constraints on DNA accessibility, through the persistence of polymerases and nascent RNAs, but also actively remodel chromatin architecture via transcription-associated changes.

Although TTF2 depletion had been previously linked to mitotic defects (Jiang et al., 2004), the mechanisms underlying these defects were not known. We reveal a minor defect in chromosome alignment, which is not related to defective KT-MT attachments and does not result in major lagging chromosomes in anaphase. Such impaired chromosome alignment, despite seemingly normal attachments, has been previously observed upon impairment of chromokinesin function (Janssen et al., 2018; Stumpff et al., 2012). However, given our results on defective chromosome assembly, we favour that the underlying hypocondensed chromatin may impact efficient alignment of chromosomes, due to potential perturbations in force-balance equilibrium (Ribeiro et al., 2009; Oliveira et al., 2005; Gerlich et al., 2006).

Indeed, we found that chromosome assembly problems are the major contributor to compromised mitotic fidelity, with anaphase bridges emerging as the most frequent defect. Similar observations were obtained in other studies, using other perturbations of mitotic transcriptional silencing (Perea-Resa et al., 2020; Sharp et al., 2020) suggesting that, independently of the specific perturbation, ongoing transcription during mitosis disrupts efficient sister chromatid resolution.

How ongoing transcription directly interferes with chromosome assembly/sister chromatid resolution remains unclear. However, our findings support a model in which alterations at the DNA substrate level—including persistent transcription-associated structures—impede efficient sister chromatid resolution. During interphase, transcription-induced topological stress is rapidly dissipated by topoisomerases, particularly DNA Topoisomerase I. In contrast, DNA Topoisomerase I is largely evicted from mitotic chromosomes (**Supplementary Fig. 8**), rendering mitotic chromatin primarily reliant on DNA Topoisomerase IIα activity. In this context, elevated transcription-associated supercoiling, potentially stabilised by R-loop accumulation, may impose persistent topological constraints. We therefore propose that transcriptional persistence into mitosis—accompanied by R-loop accumulation—functionally diverts or impedes Top2A activity at specific sites, thereby compromising complete sister chromatid resolution and giving rise to the anaphase defects observed. Although this model remains to be formally tested, it provides a plausible framework to explain the frequent DNA bridges detected despite apparently normal levels and activity of Topoisomerase IIα.

Interestingly, although we observed nascent RNAs all over mitotic chromatin, R-loop persistence is only detected at more discrete foci. These findings further suggest that specific genomic loci may be particularly prone to transcriptionally-related mitotic stress. R-loop accumulation at these sites could reflect either a causal contributor to, or a consequence of, elevated topological constraints. On one hand, defective R-loop resolution may stabilise transcription-coupled supercoiling, thereby trapping topological stress. On the other hand, impaired resolution of supercoils could hinder efficient R-loop removal, leading to their persistence into mitosis. Regardless of the directionality, these findings indicate that loci enriched in transcription-associated intermediates are especially susceptible to mitotic errors. Identifying such putative hotspots of transcription–mitosis conflict may, in the future, reveal genome regions or gene classes with increased vulnerability to chromosome segregation defects.

Recent evidence suggests that TTF2 also contributes to replisome disassembly at mitotic entry, in cells under induced replication stress (Fujisawa and Labib, 2024; Can et al., 2024). Our results, in contrast, indicate that TTF2 depletion does not result in a significant increase in RPA-coated bridges, and most UBFs and DNA bridges are indeed reverted by transcription inhibition. These findings suggest that in the absence of high replication stress, most of the segregation defects relate to transcription-associated events rather than persistent DNA replication intermediates. It is conceivable that, upon high replication stress, mitotic errors may result from a cumulative effect of transcription- and replication-related processes, both of which would simultaneously induce an increase in topological constraints and/or transcription/replication intermediates that could collectively impair proper chromosome assembly.

Altogether, our study underscores the importance of transcriptional silencing at mitotic onset for maintaining chromosome structural integrity and genome stability. Failure to properly terminate transcription leads to defective chromosome assembly, persistent topological stress, anaphase defects, and ultimately DNA damage and chromosomal instability. These observations emphasise that the coordination between transcriptional control and chromosome mechanics is central to faithful genome inheritance.

## Materials and Methods

### Cell lines and drug treatment

HeLa cells were grown in DMEM High Glucose (L0103, biowest) supplemented with 10% (v/v) Foetal Bovine Serum (FBS) (biowest) and 1% (v/v) Penicillin–Streptomycin (Gibco) at 37°C and 5% CO_2_. HeLa CENPA-mCherry cells were a gift from the Kops lab. Routine mycoplasma checks (Mycoplasmacheck Barcodes, Eurofins genomics) were conducted to ensure mycoplasma-free status. For drug treatment, cells were treated with: 5μM Triptolide (Sigma 645900) at 2h or 5h, as indicated in the figures); 1μM Mps1 inhibitor (NMSP715; Sigma Aldrich) for 20min before live-cell imaging and kept for the entire length of the imaging; 0.05μM ICRF-193 (Sigma I4659) for 2h before live-cell imaging and kept for the entire length of the imaging.

### Generation of inducible cell line expressing an siRNA-resistant form of TTF2

TTF2 human cDNA with adjacent AttB cloning sites was ordered from IDT. The target sequence of the siRNA was replaced with a synonymous DNA sequence generated with Synonymous Mutation Generator (Ong, 2021) via InFusion cloning (In-Fusion Snap Assembly MMix (Takara Bio). The wt/esiR plasmids obtained were then transferred to a destination vector (Addgene Plasmid #80474) using the Gateway™ LR Clonase™ II Enzyme mix (Invitrogen Cat. 11791020). The generated pB(TA)-NeoR-TTF2 (wt and siR) were confirmed by whole plasmid sequencing (Eurofins genomics). HeLa cells were transfected with 0.5ug of pB(TA)-NeoR-TTF2 (wt or siR) and 0.5ug of pBASE, with FUGENE HD (E2311, Promega). After 48h the media was replaced with DMEM supplemented with tet-free FBS (S181T, biowest) and Neomycin (250ug/ml). Media was replaced every other day until all cells in the untransfected control were dead. Single-cell clones were obtained by fluorescence-activated cell sorting (FACS). For wt/esiR TTF2 expression, cells were treated with doxycycline (1 μg/ml) at the time of RNAi.

TTF2 cDNA esiRNA sensitive (target sequence 2919 bp – 3320 bp) 5’-3’:

CCTTAACGGCACCTTCTTCAAGATGGAGCTTTTTGAAGGCATGCGAGAGAGCACCAAGATTTCATCTCTGTTGGCAGAATTGGAGGCAATTCAAAGAAATTCAGCATCCCAAAAGAGTGTCATTGTCTCTCAGTGGACCAACATGCTGAAAGTTGTAGCATTGCACCTGAAGAAGCATGGACTGACTTATGCCACCATCGATGGCTCTGTCAATCCCAAGCAGAGAATGGACTTGGTAGAGGCATTTAACCACTCCAGAGGCCCTCAGGTAATGCTAATCTCTCTCTTGGCCGGAGGTGTTGGTCTAAACCTGACTGGAGGAAATCACCTCTTTCTTTTGGACATGCACTGGAATCCATCACTTGAAGATCAAGCTTGTGACCGAATTTACCGAGTAGGGCA

TTF2 cDNA esiRNA resistant (target sequence 2919 bp – 3320 bp) 5’-3’:

AGCTTGAATGGAACATTTTTTAAAATGGAATTGTTCGAGGGAATGAGAGAATCCACAAAAATCAGCAGCCTCCTTGCCGAGCTTGAAGCCATCCAGCGCAACAGCGCCAGCCAGAAATCCGTGATCGTGAGCCAATGGACAAATATGCTCAAGGTGGTCGCCCTTCATCTCAAAAAACACGGCCTCACCTACGCTACAATAGACGGAAGCGTGAACCCAAAACAACGCATGGATCTTGTCGAAGCCTTCAATCATAGCCGCGGACCCCAAGTCATGTTAATAAGCCTGCTTGCTGGCGGCGTGGGCTTAAATCTCACCGGCGGCAACCATCTGTTCTTGCTTGATATGCATTGGAACCCCAGCTTGGAGGACCAGGCATGCGATAGAATCTATAGAGTCGGCCAA

### Protein knockdown via RNAi

RNAi was achieved by transfection of cells for 48h with 280 ng siRNA (EHU049891 for TTF2 and EHUFLUC as negative control, MISSION® siRNA (Sigma) using Lipofectamine RNAi MAX (Invitrogen) and Opti-MEM™ (Gibco), according to manufacturers’ protocol.

### Western Blotting

To confirm the efficient protein knockdown, cells were collected and lysed with RIPA buffer (25 mM Tris-HCl pH 7.6, 150 mM NaCl, 1% NP-40, 1% sodium deoxycholate, 0.1% SDS) supplemented with protease inhibitor (Thermo Scientific PIER78429). Samples were quantified with Bradford assay (Bio-Rad), loaded on a 4-15% gradient gel (Bio-Rad 4561084) and transferred to a nitrocellulose blot membrane. Western blot analysis was performed according to standard protocols using the following antibodies: anti-TTF2 (1:1000, Abcam ab28030) and anti-GAPDH (1:2000, Cell Signalling Technology 14C10).

### Transcripts labelling and detection

Labelling of recently transcribed RNAs (single incorporation) was performed by 5-ethynyl uridine (EU) incorporation (1 mM, 30 min in media) followed by detection via ClickIT chemistry (Click-iT® RNA Imaging Kits, Invitrogen C10329) according to manufacturer’s instructions. For sequential incorporation, cells were incubated first with 5-fluorouracil (FUrd, Thermo Scientific J62083) (1 mM, 15 min in media), washed twice with PBS and then incubated with EU (2 mM, 15 min in media). ClickIT chemistry was used for EU detection, followed by immunofluorescence for FUrd detection.

### Generation of an inducible stable cell line expressing the R-loop sensor (RHINO)

RHINO cDNA was kindly donated by Sergio Almeida lab (Martin et al., 2023) and cloned into a pcDNA5 using NotI and AlfII restriction sites. The resulting plasmid was sequenced to verify insertion and exclude any mutation (Whole plasmid sequencing, Eurofins genomics). pcDNA5-RHINO-GFP was co-transfected with pOG44 (9:1 ratio) with Lipofectamine 2000 (Thermofisher 11668027) for Flip-IN recombination into HeLa FRT/TO Trex (a gift from the Bettencourt lab). After 48h the media was replaced with DMEM supplemented with tet-free FBS (S181T, biowest) and Hygromicin (MedChem Express HY-B0490, at a concentration of 200 μg/ml). Media was replaced every other day until all cells in the untransfected control were dead. For RHINO-GFP expression, doxycycline (1 μg/ml) was added 6h previous to cell fixation with MeOH 100% at - 20 °C, Hoechst staining and slides mounting.

### Immunofluorescence and fixed-cell imaging

Cells grown on coverslips were fixed with 4% PFA in PBS for 10 min, then permeabilized with 0.5% triton in PBS for 10 min. For Top2A staining, a pre-extraction buffer was used (PHEM: 30mM HEPES, 65mM PIPES, 10mM EGTA, 2mM MgCl2, in H2O MiliQ; Adjusted to pH 6.9). For Top1, pre-extraction protocol was performed as previously described (Wiegard et al., 2021).

After blocking with 3% BSA in PBS for 30 min, cells were incubated with primary antibodies (CENPA 1:200, Invitrogen MA1-20832; BrdU, which also detects FUrd, 1:200, B2531 Merk; PICH 1:300, Abnova H00054821-M01, RPA-70 1:500, Abcam ab79398; DNA:RNA hybrids 1:200, Abcam ab234957; Mad2 1:500, Invitrogen PA5/21594; Top2A 1:1000, MBL M042-3, Top1 1:200 Abcam ab109374). After 3 washes in PBS (or PHEM), cells were stained with secondary antibodies (anti-mouse Alexa Fluor 488, anti-mouse 568, anti-rabbit Alexa 633, all used 1:1000). After wash, DNA was stained with Hoechst (1:1000), and coverslips were mounted with Vectashield (Vector H-1000; Vector Laboratories). Cells were images with a spinning disk confocal microscope (Andor Dragonfly SDC) acquired in 0.3 μm steps over a 15 μm rage.

### Live-cell imaging

HeLa cells were seeded in an 8-well Ibidi chamber (Ibidi 80826-90) and transfected as described above. 48h after transfection, cells were incubated with SiR-DNA (Spyrochrome) in media for 45 min, washed and replaced with complete media, and imaged with a spinning disk confocal microscope (Andor Dragonfly SDC) equipped with a CO_2_ (5%) and T (37 °C) controlled chamber. Three-dimensional image stacks were acquired in 0.5 μm steps over a 10 μm rage, using Olympus 40× dry objective (1.4 numerical aperture) every 3 min for 6 hours. For high temporal resolution of DMSO/ICRF-193 treated cells, imaging was performed with 1 min timelapse for 2 hours. For HeLa CENPA-mCherry imaging, cells were imaged with an Olympus 100× oil immersion objective for 4 hours, 3 min timelapse.

### Differential Retention of Topoisomerase IIα (DRT) Assay

DRT assay was performed as an optimised version of the method originally described by Agostinho *et al*. (Agostinho et al., 2004). In contrast to the original protocol, which included the catalytic inhibitor ICRF-193, only etoposide was used to avoid retention of inactive Top2A species, and etoposide concentration and incubation time was optimised to avoid saturation.

For the concentration curve (Supplementary Fig.7), HeLa wt cells were treated with etoposide for 5min (5, 25 and 50 μg/ml etop) or DMSO (as control). Immediately following the 5-minute incubation, the etoposide-containing medium was gently removed, and coverslips were placed on ice. Residual medium was removed, and approximately 70 μL of salt wash solution (300mM NaCl, 0.5% TritonX-100, 1mM PMFS, 1x Anti-proteases cocktail, in PHEM buffer) was added to each coverslip, incubated on ice for 1 min 30 s. The salt wash solution was removed, and cells were fixed by adding 4% PFA, and incubated for 10 min at room temperature. After fixation, PFA was removed and cells were incubated in PHEM buffer containing 0.5% Triton X-100, for 5 min to permeabilize cellular membranes. Finally, the PHEM–Triton solution was removed, cells were washed with PBS, and samples were ready for downstream immunofluorescence (IF).

Cells were transfected with siRNAs targeting TTF2 or CTRL control as described previously (see Protein knockdown via RNAi section). Transfections were carried out in standard culture medium, and cells were allowed to incubate for 48 h to achieve efficient knockdown. After 48 h, cells were treated with 12.5 μg/mL etoposide for 5 min. After 1 min 30 s salt wash on ice, cells were fixed by adding ice-cold 100% methanol, and incubated for 15 min at -20 °C. After fixation, methanol was removed, and cells were prepared as described above.

### Fluorescence *In Situ* Hybridisation (FISH)

Cells grown on glass coverslips were fixed in cold MeOH:Acetic Acid (3:1) for 10 min. Coverslips were treated with EtOH series (75, 85, 100%) for 3 min each and let dry. Whole chromosome paint probes (XCP 17-FITC, whole chr 17 FISH; XCP 12 Orange, whole chr 12 FISH MetaSystems) were added on coverslips, covered and denatured at 75 C for 2 min, followed by an hybridization step at 37 °C overnight in a humid chamber. The following day, coverslips were washed 4 min with 0.4 SSX at 74 °C followed by 2 washes in 2XSSC 0.05 % tween at room temperature. Finally, coverslips were rinsed in dH_2_O, DNA was stained with Hoechst, and coverslips mounted on glass slide with Vectashield.

### Imaging analysis

Image analysis was performed using Fiji software.

For EU retention, images were analysed after max-intensity z-projection. DNA and whole cell areas were selected with automatic and manual threshold, respectively, on Hoechst channel, and the mean intensity of EU measured. To obtain the transcripts retention values (Figure 1 and Suppl. Figure 2), the mean intensity over the DNA area was divided by the mean intensity over the whole cell area. Analysis of FUrd/EU retention was performed manually on blinded images; cells were scored based on the presence of each single staining or the combination of the two.

For quantification of Top2A (Figure 4) and EU on TRP treated cells (Supplementary Fig.5), cells were selected with automatic threshold on Hoechst channel and the mean intensity measured.

Metaphase width measurement in live-cell imaging was performed in metaphase one frame before AO (-3 min). A line perpendicular to the metaphase plate was drawn, and measurements of the DNA occupied underneath were taken.

For centromere alignment analysis in metaphase (Figure2) CENPA-mCherry area was measured after max-intensity z-projection of live-cell imaging, at each indicated time points before AO. Images were first smoothed, and metaphase plate was selected based on SiR-DNA threshold. The CEN signal was then thresholded manually to create a binary mask. The mask was converted to points, a convex hull was generated, and the outline was smoothed by spline fitting. The resulting ROI was measured to calculate the total area occupied by CEN.

Inter-centromeric distance (Supplementary Fig.4) was analysed on CENPA-mCherry live-cell imaging through each z acquired. The distance between 2 adjacent CENPA dots was measured one frame before AO (-3 min).

For single chromosomes volumes on FISH images, a custom Fiji macro was used to measure 3D volumes of segmented objects. The macro processes single files and uses pre-saved ROIs drawn around each single chromosomes. For each ROI, the macro extracts the corresponding channel, thresholds the signal to create a binary mask and applies 3D convex hull analysis (3D_Convex_Hull plugin). The resulting object and convex hull volumes were saved.

Mitotic errors and MN enrichment were blindly evaluated based on DNA staining in live (SiR-DNA) and fixed cells (Hoechst) or based on the presence of UFBs. Cells where segregation fidelity could not be assigned were excluded from the analysis.

For DNA:RNA hybrids analysis (Figure 5F), S9.6 antibody foci number was assessed by using the Find Maxima plugin with prominence 250 on a selected max projection encompassing the entire DNA in metaphase.

For R-loops analysis of RHINO-GFP anaphase/telophase errors, cells were grouped based on the presence (faulty mitoses) or absence (error-free mitoses) of mitotic errors. Cells were then analysed on each single Z plane and, when erroneous, it was assessed whether the RHINO-GFP signal was involved in the error.

## Statistical analysis

All graphs were prepared, and statistical testing performed in the Prism software (version 10, GraphPad) using α=0.05. Details of the statistical methods applied are included in the figure legends.

## Acknowledgements

We thank all members of the Oliveira lab for fruitful discussions. We thank the Imaging Facility of Gulbenkian Institute for Molecular Sciences (GIMM). Special thanks to Carolina Pereira for macro design, Geert Kops (Hubrecht Institute) for mCherry-CENPA cell line, Sérgio Almeida (GIMM) for RHINO constructs/sequences and Monica Bettencourt (GIMM) for HeLa FRT/TO Trex cell line.

## References

Agostinho, M., Rino, J., Braga, J., Ferreira, F., Steffensen, S., & Ferreira, J. (2004). Human Topoisomerase II: Targeting to Subchromosomal Sites of Activity during Interphase and Mitosis. Molecular Biology of the Cell, 15, 2388–2400. 10.1091/mbc.E03-08

Biebricher, A., Hirano, S., Enzlin, J. H., Wiechens, N., Streicher, W. W., Huttner, D., Wang, L. H. C., Nigg, E. A., Owen-Hughes, T., Liu, Y., Peterman, E., Wuite, G. J. L., & Hickson, I. D. (2013). PICH: A DNA Translocase Specially Adapted for Processing Anaphase Bridge DNA. Molecular Cell, 51(5), 691–701. 10.1016/j.molcel.2013.07.016

Blower, M. D. (2016). Centromeric Transcription Regulates Aurora-B Localization and Activation. Cell Reports, 15(8), 1624–1633. 10.1016/j.celrep.2016.04.054

Bou-Nader, C., Bothra, A., Garboczi, D. N., Leppla, S. H., & Zhang, J. (2022). Structural basis of R-loop recognition by the S9.6 monoclonal antibody. Nature Communications, 13(1). 10.1038/s41467-022-29187-7

Can, G., Shyian, M., Krishnamoorthy, A., Lim, Y., Wu, R. A., Zaher, M. S., Raschle, M., Walter, J. C., & Pellman, D. S. (2024). TTF2 promotes replisome eviction from stalled forks in mitosis. 10.1101/2024.11.30.626186

Carmo, C., Coelho, J., Silva, R. D., Tavares, A., Boavida, A., Gaetani, P., Guilgur, L. G., Martinho, R. G., & Oliveira, R. A. (2023). A dual-function SNF2 protein drives chromatid resolution and nascent transcript removal in mitosis. EMBO Reports, 24(9). 10.15252/embr.202256463

Chan, Y. W., Fugger, K., & West, S. C. (2018). Unresolved recombination intermediates lead to ultra-fine anaphase bridges, chromosome breaks and aberrations. Nature Cell Biology, 20(1), 92–103. 10.1038/s41556-017-0011-1

Chanboonyasitt, P., & Chan, Y. W. (2021). Regulation of mitotic chromosome architecture and resolution of ultrafine anaphase bridges by PICH. Cell Cycle (Vol. 20, Number 20, pp. 2077–2090). Taylor and Francis Ltd. 10.1080/15384101.2021.1970877

Contreras, A., & Perea-Resa, C. (2024). Transcriptional repression across mitosis: mechanisms and functions. Biochemical Society Transactions (Vol. 52, Number 1, pp. 455–464). Portland Press Ltd. 10.1042/BST20231071

Finardi, A., Massari, L. F., & Visintin, R. (2020). Anaphase bridges: Not all natural fibers are healthy. In Genes (Vol. 11, Number 8, pp. 1–29). MDPI AG. 10.3390/genes11080902

Fujisawa, R., & Labib, K. P. M. (2024). TTF2 drives mitotic replisome disassembly and MiDAS by coupling the TRAIP ubiquitin ligase to Polε. 10.1101/2024.12.01.626218

Gerlich, D., Hirota, T., Koch, B., Peters, J. M., & Ellenberg, J. (2006). Condensin I stabilizes chromosomes mechanically through a dynamic interaction in live cells. Current Biology, 16(4), 333–344. 10.1016/j.cub.2005.12.040

Huang, K. C., Gao, H., Yamasaki, E. F., Grabowski, D. R., Liu, S., Shen, L. L., Chan, K. K., Ganapathi, R., & Snapka, R. M. (2001). Topoisomerase II Poisoning by ICRF-193. Journal of Biological Chemistry, 276(48), 44488–44494. 10.1074/jbc.M104383200

Janssen, L. M. E., Averink, T. V., Blomen, V. A., Brummelkamp, T. R., Medema, R. H., & Raaijmakers, J. A. (2018). Loss of Kif18A Results in Spindle Assembly Checkpoint Activation at Microtubule-Attached Kinetochores. Current Biology, 28(17), 2685–2696.e4. 10.1016/j.cub.2018.06.026

Jennifer C Waters, Robert V Skibbens, & E D Salmon. (1996). Oscillating mitotic newt lung cell kinetochores are, on average, under tension and rarely push. Journal of Cell Science.

Jiang, Y., Liu, M., Spencer, C. A., & Price, D. H. (2004). Involvement of transcription termination factor 2 in mitotic repression of transcription elongation. Molecular Cell, 14(3), 375–386. 10.1016/S1097-2765(04)00234-5

Lebreton, J., Colin, L., Chatre, E., & Bernard, P. (2024). RNAP II antagonizes mitotic chromatin folding and chromosome segregation by condensin. Cell Reports, 43(3). 10.1016/j.celrep.2024.113901

Liu, M., Xie, Z., & Price, D. H. (1998). A human RNA polymerase II transcription termination factor is a SWI2/SNF2 family member. Journal of Biological Chemistry, 273(40), 25541–25544. 10.1074/jbc.273.40.25541

Martin, R. M., De Almeida, M. R., Gameiro, E., & De Almeida, S. F. (2023). Live-cell imaging unveils distinct R-loop populations with heterogeneous dynamics. Nucleic Acids Research, 51(20), 11010–11023. 10.1093/nar/gkad812

McNulty, S. M., Sullivan, L. L., & Sullivan, B. A. (2017). Human Centromeres Produce Chromosome-Specific and Array-Specific Alpha Satellite Transcripts that Are Complexed with CENP-A and CENP-C. Developmental Cell, 42(3), 226–240.e6. 10.1016/j.devcel.2017.07.001

Montecucco, A., Zanetta, F., & Biamonti, G. (2015). Molecular mechanisms of etoposide. In EXCLI Journal (Vol. 14, pp. 95–108). Leibniz Research Centre for Working Environment and Human Factors. 10.17179/excli2014-561

Oliveira, R. A., Coelho, P. A., & Sunkel, C. E. (2005). The Condensin I Subunit Barren/CAP-H Is Essential for the Structural Integrity of Centromeric Heterochromatin during Mitosis. Molecular and Cellular Biology, 25(20), 8971–8984. 10.1128/mcb.25.20.8971-8984.2005

Oliveira, R. A., Hamilton, R. S., Pauli, A., Davis, I., & Nasmyth, K. (2010). Cohesin cleavage and Cdk inhibition trigger formation of daughter nuclei. Nature Cell Biology, 12(2). 10.1038/ncb2018

Ong, J. Y. (2021). Synonymous Mutation Generator: a web tool for designing RNAi-resistant sequences. 10.1101/2021.01.02.425100

Palozola, K. C., Liu, H., Nicetto, D., & Zaret, K. S. (2017). Low-Level, Global Transcription during Mitosis and Dynamic Gene Reactivation during Mitotic Exit. Cold Spring Harbor Symposia on Quantitative Biology, 82, 197–205. 10.1101/sqb.2017.82.034280

Pelham-Webb, B., Polyzos, A., Wojenski, L., Kloetgen, A., Li, J., Di Giammartino, D. C., Sakellaropoulos, T., Tsirigos, A., Core, L., & Apostolou, E. (2021). H3K27ac bookmarking promotes rapid post-mitotic activation of the pluripotent stem cell program without impacting 3D chromatin reorganization. Molecular Cell, 81(8), 1732–1748.e8. 10.1016/j.molcel.2021.02.032

Perea-Resa, C., Bury, L., Cheeseman, I. M., & Blower, M. D. (2020). Cohesin Removal Reprograms Gene Expression upon Mitotic Entry. Molecular Cell, 78(1), 127–140.e7. 10.1016/j.molcel.2020.01.023

Ramos-Alonso, L., Holland, P., Le Gras, S., Zhao, X., Jost, B., Bjørås, M., Barral, Y., Enserink, J. M., & Chymkowitch, P. (2023). Mitotic chromosome condensation resets chromatin to safeguard transcriptional homeostasis during interphase. Proceedings of the National Academy of Sciences of the United States of America, 120(4). 10.1073/pnas.2210593120

Ribeiro, S. A., Gatlin, J. C., Dong, Y., Joglekar, A., Cameron, L., Hudson, D. F., Farr, C. J., Mcewen, B. F., Salmon, E. D., Earnshaw, W. C., & Vagnarelli, P. (2009). Condensin Regulates the Stiffness of Vertebrate Centromeres. Molecular Biology of the Cell, 20. 10.1091/mbc.E08

Sarlós, K., Biebricher, A. S., Bizard, A. H., Bakx, J. A. M., Ferreté-Bonastre, A. G., Modesti, M., Paramasivam, M., Yao, Q., Peterman, E. J. G., Wuite, G. J. L., & Hickson, I. D. (2018). Reconstitution of anaphase DNA bridge recognition and disjunction. Nature Structural and Molecular Biology, 25(9), 868–876. 10.1038/s41594-018-0123-8

Sharp, J. A., Perea-Resa, C., Wang, W., & Blower, M. D. (2020). Cell division requires RNA eviction from condensing chromosomes. Journal of Cell Biology, 219(11). 10.1083/JCB.201910148

Stumpff, J., Wagenbach, M., Franck, A., Asbury, C. L., & Wordeman, L. (2012). Kif18A and Chromokinesins Confine Centromere Movements via Microtubule Growth Suppression and Spatial Control of Kinetochore Tension. Developmental Cell, 22(5), 1017–1029. 10.1016/j.devcel.2012.02.013

Titov, D. V., Gilman, B., He, Q. L., Bhat, S., Low, W. K., Dang, Y., Smeaton, M., Demain, A. L., Miller, P. S., Kugel, J. F., Goodrich, J. A., & Liu, J. O. (2011). XPB, a subunit of TFIIH, is a target of the natural product triptolide. Nature Chemical Biology, 7(3), 182–188. 10.1038/nchembio.522

Wiegard, A., Kuzin, V., Cameron, D. P., Kouzine, F., Natsume, T., & Baranello, L. (2021). Topoisomerase 1 activity during mitotic transcription favors the transition from mitosis to G1. Molecular Cell, 81, 1–18. 10.1016/j.molcel.2021.10.015

Wojenski, L. A., Wainman, L. M., Villafano, G., Kuhlberg, C., Nanda, P., & Core, L. (2021). Multiple modes of regulation control dynamic transcription patterns during the mitosis-G1 transition *bioRxiv* 2021.06.22.449286; doi: 10.1101/2021.06.22.449286.

