## Supplementary Fig. for "Removal of nascent transcripts by TTF2 is required for sister chromatid resolution in human cells"

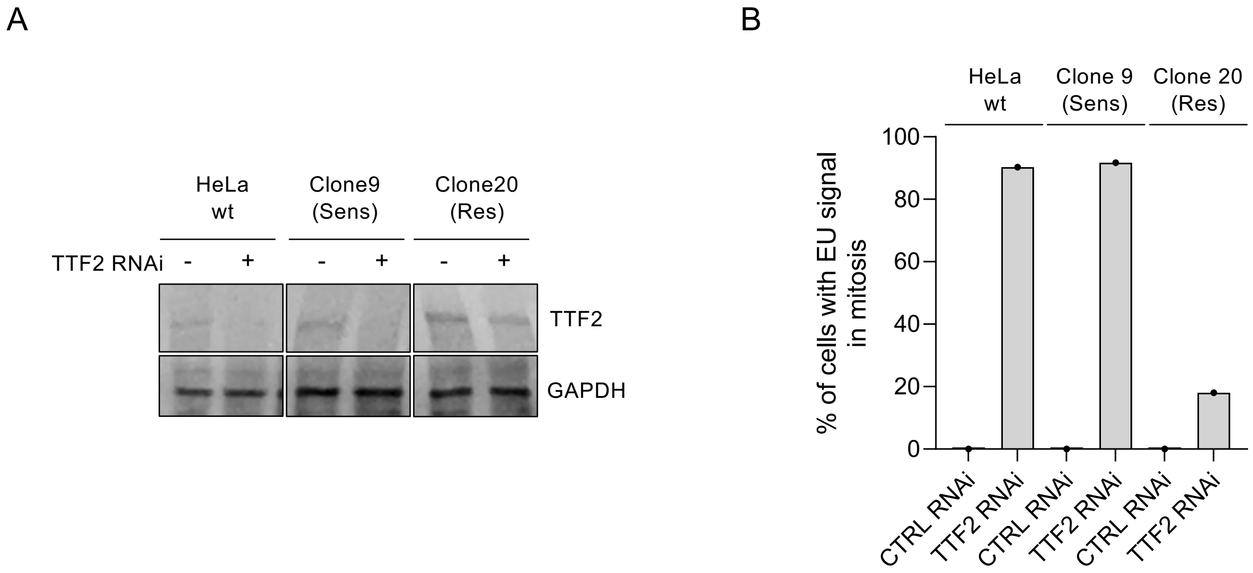


**Supplementary Figure 1. EU retention in metaphase is caused by TTF2 KD and not by off-target events**

**A**) Western blot of parental HeLa cell and the selected siRNA resistant/sensitive clones upon CTRL/TTF2 RNAi. Top band, TTF2 (130 KDa), bottom band GAPDH (39 KDa). **B**) Quantification of EU+ metaphase cells upon TTF2 KD in HeLa wt and in the selected siRNA sensitive or resistant clones. Bars indicate the mean, data from 1 experiment, where at least 30 mitotic cells per condition were analysed.

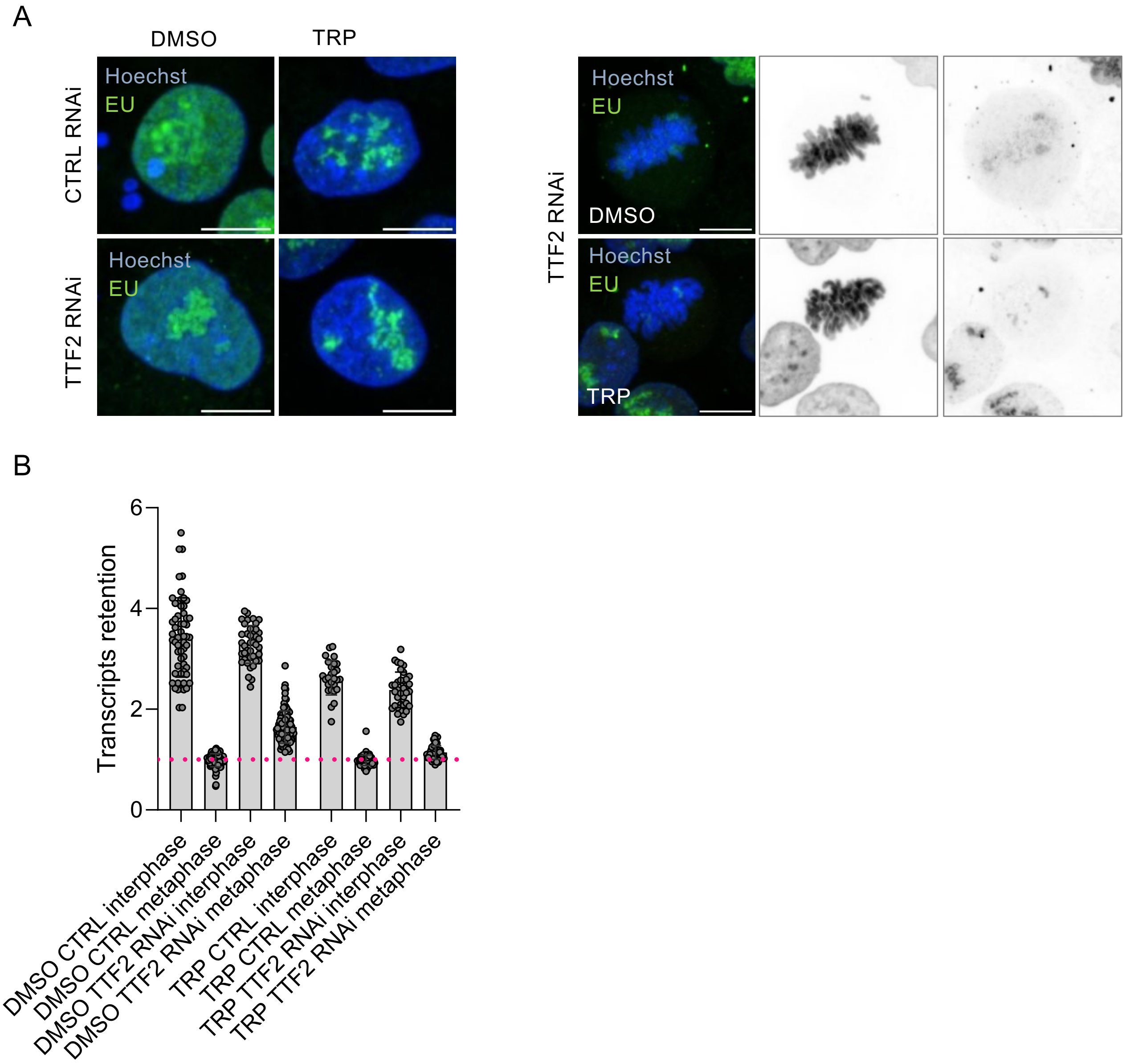


**Supplementary Figure 2. EU transcripts retention in metaphase is reverted upon triptolide (TRP) treatment**
**A**) Representative images of HeLa cells upon CTRL/TTF2 RNAi and EU incorporation after 5h TRP treatment. Interphase cells on the left side of the panel, metaphase cells on the right. DNA is in blue (Hoechst) and nascent transcripts are in green (EU). **B**) EU signal retention on DNA area relative to whole cell area in interphase and in metaphase in DMSO and TRP treated cells. Each dot represents one cell, with bars indicating the mean value of 3 independent experiments. Pink dotted line indicate the baseline for no chromatin enrichment of the EU signal (=1). Scale bars are all 10 um.

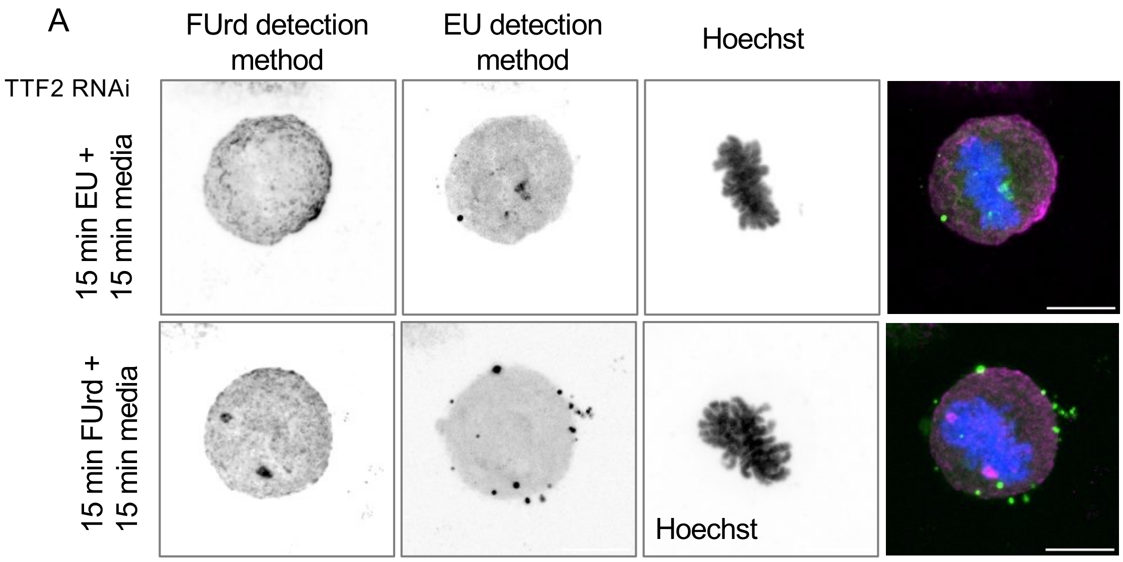


**Supplementary Figure 3. FUrd and EU analogues cannot be recognized by each other's detection method**
**A**) Representative images of HeLa metaphase cells upon TTF2 RNAi and EU or FUrd incorporation followed by 15 min in standard media in the absence of analogue. EU is in green, FUrd is in magenta, DNA in blue. Scale bars are all 10 um.

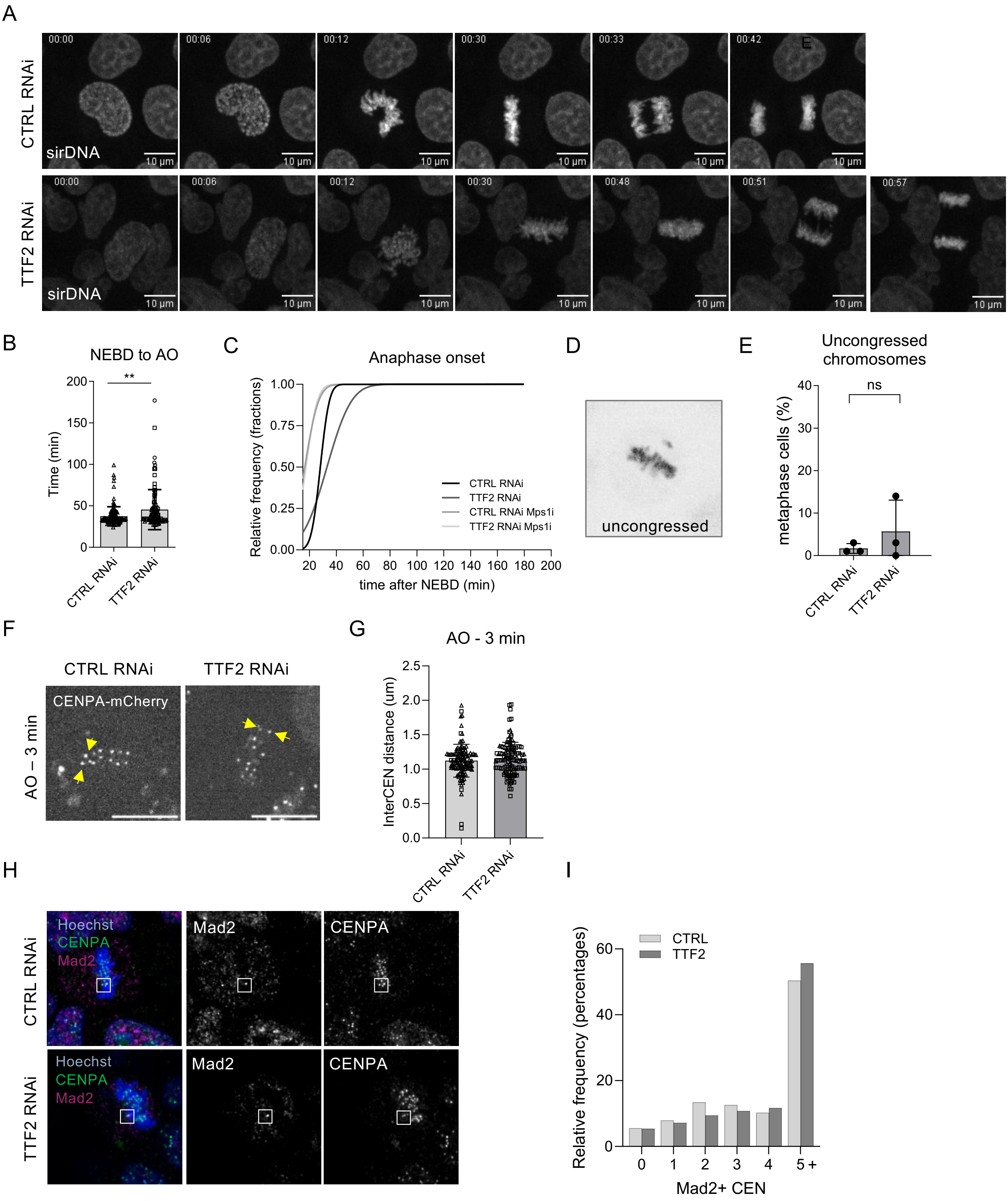


**Supplementary Figure 4. TTF2-depleted cells do not show evidence of the canonical triggers for SAC activation**

**A**) Representative snap-shots from live-cell imaging of HeLa mitotic progression upon CTRL/TTF2 RNAi. **B**) Measurements of time spent in mitosis (time from NEBD to AO) from live-cell imaging of CTRL/TTF2 KD cells. Each dot represents one cell, bars indicating the mean and SD from 3 independent experiments. **p= 0.0012. **C**) Cumulative frequency (fraction of all) cells entering anaphase in CTRL/TTF2 cells. Data are from 3 independent experiments, 1 experiment for Mps1i treated cells. **D**) Example of uncongressed chromosome in metaphase. **E**) Quantification of cells with uncongressed chromosomes (%), defined as chromosomes not aligned at metaphase plate formation. Bars indicate the mean of 3 independent experiments. **F**) Representative images of CENPA-mCherry signal in HeLa metaphase cells one time-frame before AO (-3 min). Yellow arrows indicate pairs of centromere. **G**) Measurements of inter-centromere distance of pair of centromeres as indicated in F. Each dot indicate one pair of signal at the centromere, bars indicate the mean and SD of 3 independent experiments. **H**) Example images of metaphase cells with centromeric staining (CENPA, green) and Mad2 (magenta ).**I**) Percentage of prometaphase/metaphase cells with Mad2+ signal at centromere, data from 1 experiment. *p*-values were calculated using a Student's t-test. Scale bars are all 10 um.

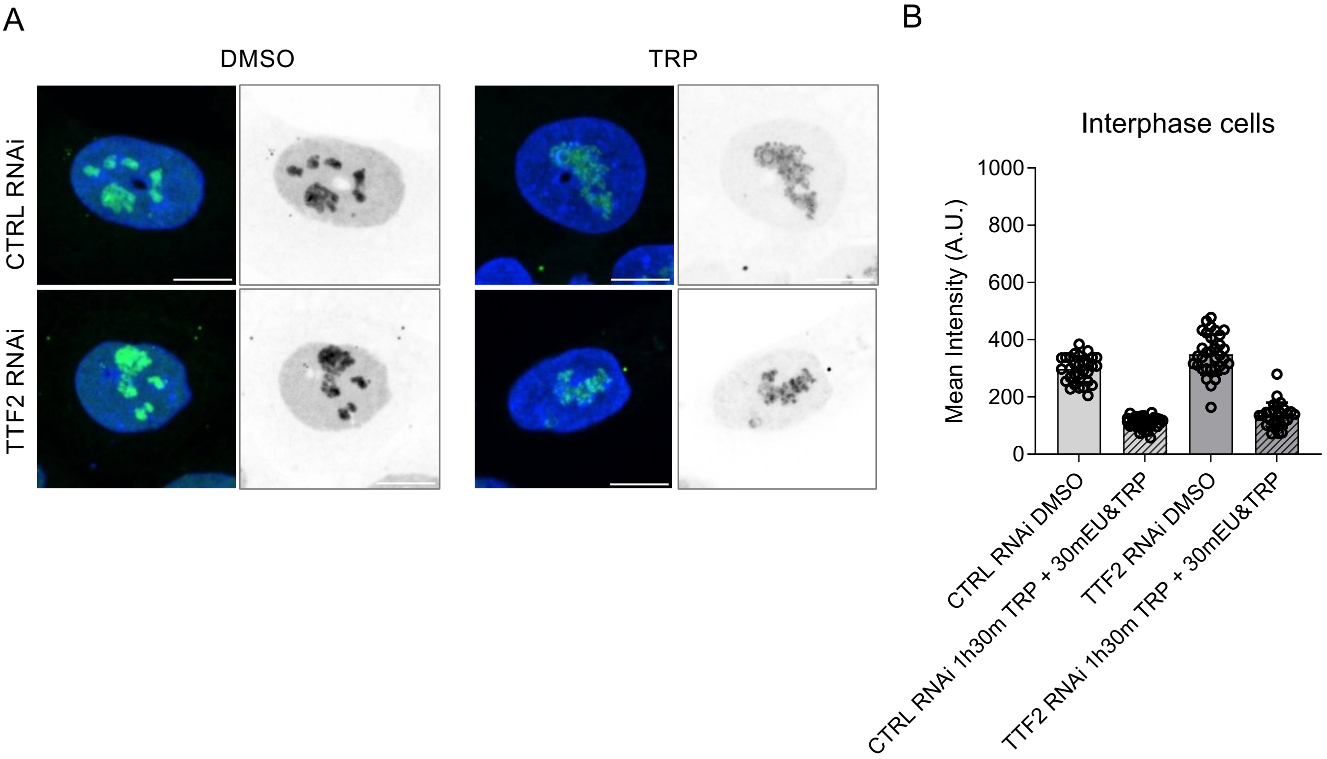


**Supplementary Figure 5. Short triptolide treatment (2h) is sufficient to remove most of the transcripts from chromatin**
**A)** Representative images of HeLa cells upon TTF2 RNAi and TRP treatment (1h + 30 min with EU or 1h30 min +30 min EU). B) Quantification of EU signal (mean intensity over DNA area) in interphase cells treated with DMSO or TRP as indicated. Each dot represents one cell, bars indicate the mean. Data from 1 experiment, where at least 30 cells were analysed.

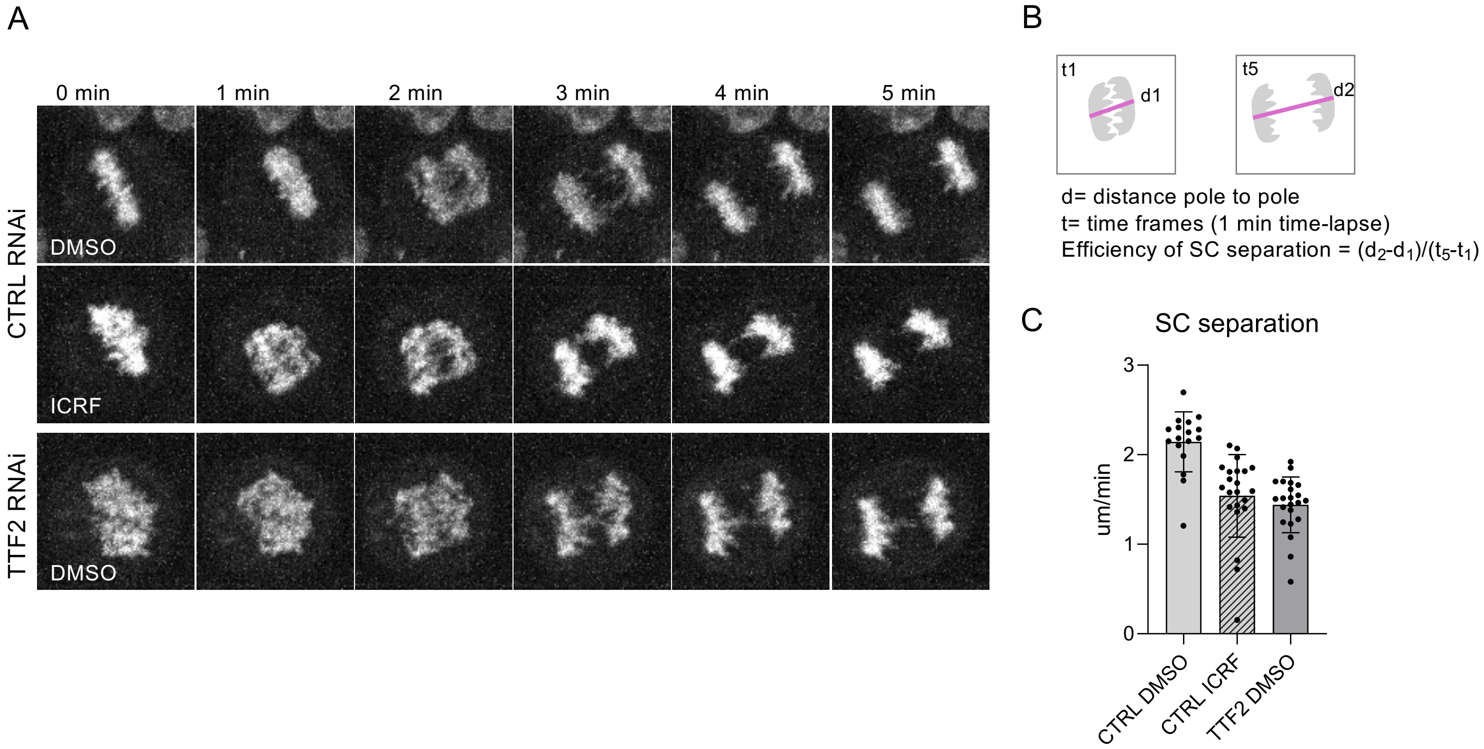


**Supplementary Figure 6. Impaired sister chromatid resolution upon TTF2 loss**

**A**) Representative snap-shots from live-cell imaging of HeLa mitotic progression upon CTRL/TTF2 RNAi and ICRF treated cells at higher temporal resolution (1 min/frame). **B**) Schematic of the measurements to assess the efficiency of SC separation **C**) Quantification of the time needed for sister chromatids separation, measured as the distance (um) travelled per minute. Each dot represents one cell, data from 1 experiment.

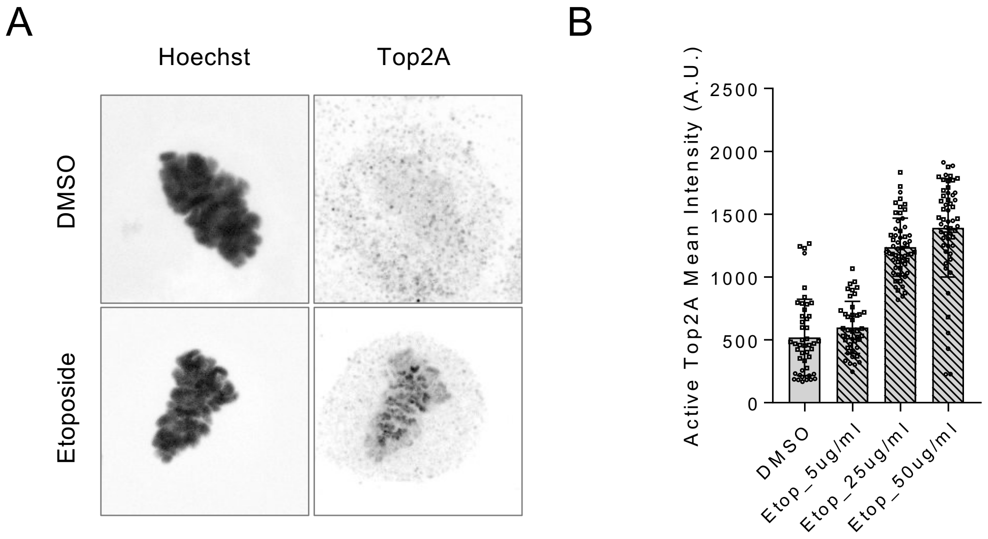


**Supplementary Figure 7**. **Retention of active Top2A at chromatin after etoposide treatment in HeLa wt cells A)** Representative images of HeLa cells upon DMSO or etoposide treatment (5 minutes treatment with 50ug/ml etoposide). **B)** Quantification of active Top2A signal (mean intensity) on chromatin of mitotic cells treated with DMSO or different concentrations of etoposide (Etop) as indicated.

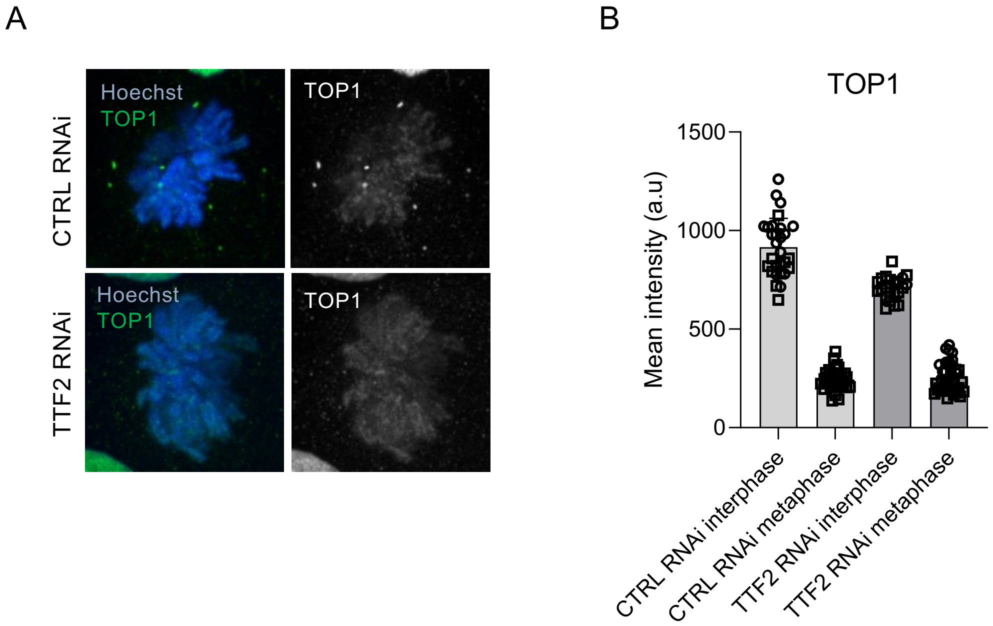


**Supplementary Figure 8. Top1 is efficiently removed even in conditions of RNA pol II retention on mitotic chromatin**
**A**) Representative images of Hela CTRL/TTF2 KD cells stained with Top1 (green), upon pre-extraction. **B**) Quantification of mean intensity values of Top1 on chromatin in interphase and metaphase in CTRL/TTF2 KD cells. Each dots represent one cell, bars indicate mean and SD, data comes from 2 independent experiments.
